# The collision of two genomes threatens global food security

**DOI:** 10.64898/2025.12.25.696198

**Authors:** Henry L. North, Gabriela Montejo-Kovacevich, Douglas Amado, Ian A. Warren, Marek Kucka, Abby Williams, Yingguang Frank Chan, Tom Walsh, Alberto Soares Corrêa, Celso Omoto, Chris D. Jiggins

## Abstract

Human activity alters selection pressures and redistributes biodiversity, creating opportunities for hybridization when different species come into secondary contact. It is now well-established that ancient hybridization plays an important role in adaptive radiation ^1^, but its role in recent anthropogenic adaptation remains unclear. We investigated this in a study of hybridizing native and invasive *Helicoverpa* moths in Brazil, which are among the most economically damaging crop pests globally. Native *H. zea* has recently evolved Bt resistance and is better adapted to maize as a host plant, while invasive *H. armigera* is resistant to pyrethroid pesticides and is better adapted to soybean. Using a decade-long time series of 975 genomes from both species, we demonstrate rapid, bidirectional adaptive introgression of structural variants that cause resistance to two distinct pesticides. In one direction, a pyrethroid resistance gene from *H. armigera* (*CYP337B3*) is now almost fixed in native *H. zea*. In the other direction, ~30% of recently-sampled invasive *H. armigera* carry a *H. zea*-derived trypsin repeat cluster. Using controlled crosses, we show that this trypsin repeat cluster confers resistance to Bt (Cry1Ac) soy in the *H. armigera* genomic background. Thus, hybridization has combined *H. armigera* adaptation to soy with H. zea Cry1Ac resistance to facilitate exploitation of Brazilian Bt soy monoculture — a vast anthropogenic habitat exceeding the area of Germany. Our results show that the combinatorial mechanisms that produce biodiversity can also drive rapid anthropogenic adaptation on ecological timescales, with consequences for global food security.

## Introduction

Human activity imposes natural selection at unprecedented scales through changes in land use, climate, and chemical exposure. At the same time, humans are redistributing biodiversity across the globe, resulting in the introduction of non-native species to novel ecological conditions. As a consequence, biological invasion events — where ecological opportunists establish and spread in new environments — are occurring at an accelerating rate ^2^. As species’ distributions shift, once-isolated lineages can come into secondary contact and may hybridize. This confluence of strong natural selection and admixture can create conditions for evolutionary rescue, with far-reaching implications from conservation to food security ^3,4^.

It is now well-established that introgression between previously isolated lineages can produce novel hybrid trait combinations that drive adaptive diversification over thousands or millions of years ^1^. Central to this ‘combinatorial mechanism’ is the amalgamation of modular genetic architectures into new configurations that facilitate the occupation of new niche space. Since admixture introduces adaptive variants at high frequency, which have already been optimised by selection in parental linages, adaptation can proceed rapidly. For example, hybridization brought together two wing colour developmental switches selected in independent *Heliconius* butterfly linages, resulting in a novel recombinant phenotype underlying adaptation and reproductive isolation ^5^. Similarly, novel traits required for ecological divergence in *Helianthus* sunflowers resulted from a combination of alleles that arose in distinct ancestral lineages ^6^. A handful of studies have shown that anthropogenic activity can lead to unidirectional adaptive introgression ^7–10^. Given the role of combinatorial evolution in ancient adaptive diversification, this begs the question: could similar mechanisms drive adaptation at even shorter timescales relevant to anthropogenic change? Cases of secondary contact between native and invasive species are particularly tractable systems in which to investigate this question.

Secondary contact between previously isolated noctuid moth species of the genus *Helicoverpa* present a remarkable natural experiment in rapid adaptation. In 2013 *Helicoverpa armigera* was detected as an invasive species in Brazil, where it encountered its sister species, *H. zea*. While *H. armigera* has a broad distribution spanning most of the eastern hemisphere, *H. zea* — which diverged in allopatry 1.5-2MYA — is native to the Americas ^11^. The two species are among the world’s most destructive agricultural pests, and the establishment of *H. armigera* in South America resulted in annual economic damage of around $US 10^7^ ^12^. Highly polyphagous larvae feed on the reproductive structures of major crops including soy, maize, and cotton ^13^. The species’ broad-scale pesticide resistance, short generation times (up to 11/yr), high fecundity (>1,000 eggs/female), and long-distance dispersal (up to 40 km/night) enable explosive population growth ^14–17^. Despite being difficult to distinguish morphologically, the two species are separated by strong prezygotic behavioural and mechanical barriers, which makes them difficult to cross in captivity ^18^. It was therefore a surprise when genomic data revealed occasional interspecific hybridization in Brazil ^19,20^. This resulted in adaptive introgression of the *de novo* chimeric pyrethroid pesticide resistance gene *CYP337B3* from invasive *H. armigera* to native *H. zea* populations in Brazil and subsequently North America ^21,22^. Previous work hinted at a single bout of locus-specific introgression ^21^. However, the two species display complementary ecological adaptations, leading to speculation that further introgression could drive ongoing adaptation to control measures ^20,23^.

Invasive *H. armigera* primarily affects soybean and cotton — the former being the most abundant crop in Brazil, comprising 47.3×10^6^ hectares in the 2024-25 crop season, an area exceeding the total size of Germany ^24,25^. However, following its outbreak in 2013, *H. armigera* damage to cotton and soy has been effectively suppressed through the widespread use of soy crops genetically modified to express *Bacillus thuringiensis* (Bt) toxins such as Cry1Ac ^26,27^. In contrast, native *H. zea* cannot complete its lifecycle on soy; it primarily affects the second most-abundant Brazilian crop — maize — and is tolerant to Cry toxins, which have been expressed in North American maize and cotton since the late 1990s ^28^. The coexistence of these species raises the question of whether hybridization could combine the Bt tolerance of native *H. zea* with the soy adaptation of invasive *H. armigera* to facilitate adaptation to Bt soybean monoculture in Brazil.

Here, we study the recent adaptive consequences of hybridization using a decade-long whole-genome time series of 975 field-collected moths from across Brazil, combined with the genomes of a further 149 individual offspring from a cross between Bt-resistant and susceptible *H. armigera*. This unique dataset captures the complete time course of the invasion and its aftermath. We reveal ongoing hybridisation and bidirectional gene flow, identify a trypsin gene cluster introgressed from *H. zea* into *H. armigera*, and show experimentally that it confers resistance to Cry1Ac-expressing soy in the *H. armigera* genomic background. These architectures are produced by the sorting of alleles at major-effect loci independently optimised by anthropogenic selection in native and invasive pests. These results provide a rare insight into rapid adaptation over ecological timescales through selection on novel genomic architectures for pesticide resistance.

## Results

### Outcome of secondary contact

We first sought to characterise the outcome of secondary contact between these species over time by quantifying individual-level ancestry across the full set of Brazilian *Helicoverpa* samples (*n*=975, Extended Data Table 1). There are various possible outcomes to hybridization. At one extreme, the two species could collapse into a single hybrid swarm, predicting a pattern of increased intermediate-ancestry individuals over time. At the other extreme, reinforcing selection would predict declining intermediate-ancestry individuals and declining ancestry heterozygosity over time. To distinguish between these patterns, we used non-admixed samples of both species to assign ancestry (Fig. 1A)

**Figure 1.**
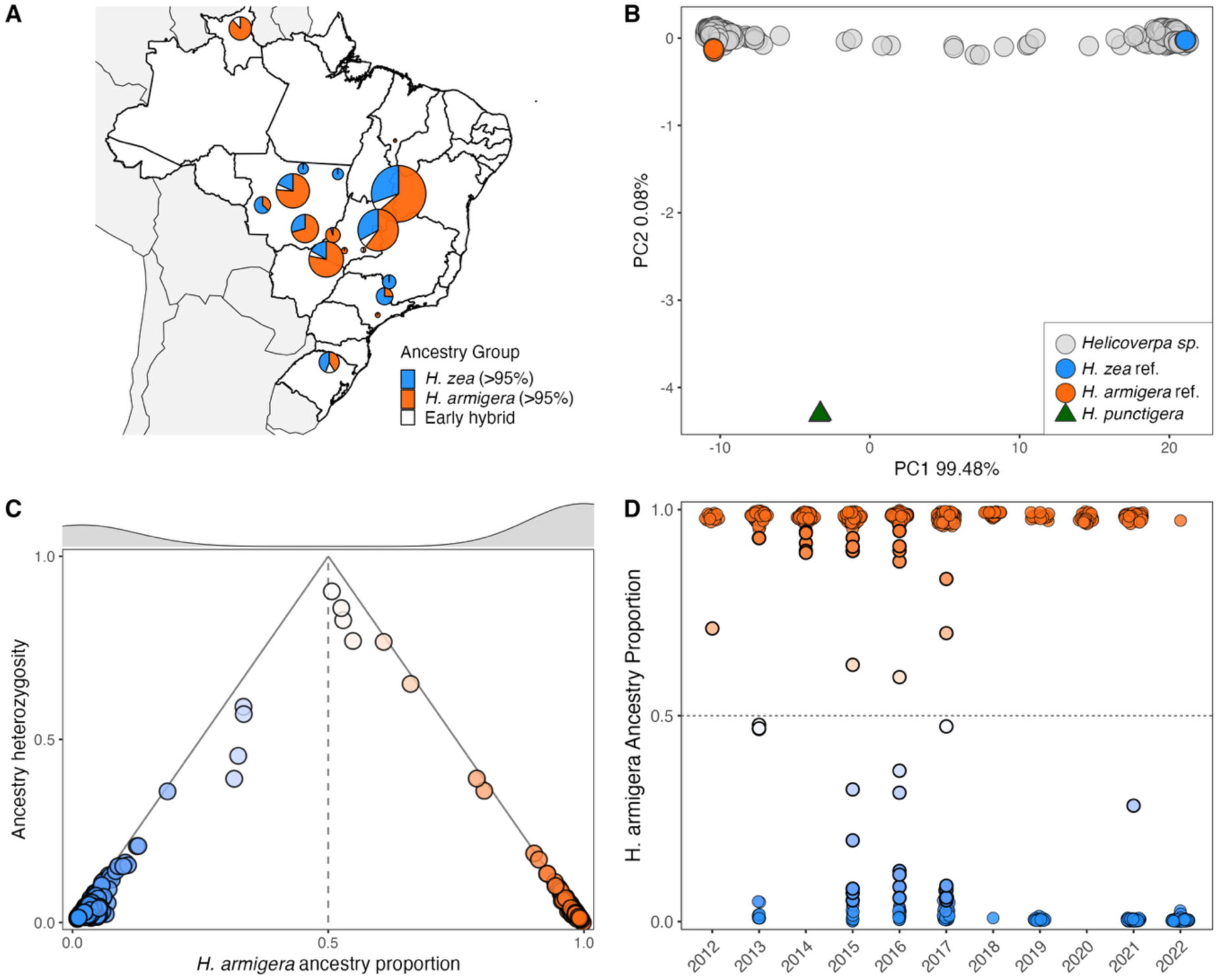
**(A)** Sample sites of individual genomes across all years (*N*=975). Circle size corresponds to per-site sample size, colour corresponds to ancestry group. **(B)** Principal components analysis (PCA) of reference non-admixed samples and Brazilian samples. **(C)** Ancestry proportion and heterozygosity of Brazilian samples in Call Set 2. **(D)** *H. armigera* ancestry proportion of Brazilian samples in the genotype likelihood dataset for each year.

Most Brazilian samples clustered with reference *H. armigera* and *H. zea* samples in principal component space, with a minority of intermediate individuals (Fig. 1B). Ancestry proportions estimated using diagnostic loci (Fig. 1C) and genotype likelihoods (Fig. 1D, Extended Data Fig 1) confirm that most samples primarily carry ancestry from one species or the other. Early-generation hybrids were rare. Nevertheless, the vast majority of samples were advanced-generation back-crosses carrying a small proportion of ancestry from the alternative species. Patterns of ancestry across space and time indicate an established mosaic hybrid zone throughout Brazil (Extended Data Figs. 2A,B) consistent with previous reports ^29^. The dominant ancestry of samples was predicted by the crop on which they were sampled: *H. armigera* on soy and cotton, and *H. zea* on maize (Extended Data Fig. 3). Proportionally more early-generation hybrids (those with <95% ancestry of either species) were found where the two species overlapped spatially and temporarily (Extended Data Fig. 2C).

The frequency of advanced-generation back-crosses in early samples, combined with the lack of spatial or temporal change in ancestry, confirms that *H. armigera* had established in South America well before its official discovery ^30^. Although the ancestry average heterozygosity decreased over time, early-generation hybrids were still found in recent years, indicating rare but ongoing hybridization (Fig. 1D, Extended Data Fig. S4). In summary, the data show stable coexistence of the two species, but with extensive interspecific gene flow: most individuals carry a small proportion of alleles from the alternate species. We next sought to identify those introgressed alleles.

### Genome-wide patterns of introgression

To identify loci with disproportionate representation of the alternate species’ ancestry, we assigned individuals to species using an ancestry proportion threshold of 0.5 (Fig. 1C) and measured excess allele sharing with reference non-admixed heterospecific samples using the statistic *f_d_*. We used two test topologies, which allowed us to distinguish between introgression from South American *H. armigera* to South American *H. zea* and introgression in the opposite direction (see Methods). Readers familiar with earlier *Helicoverpa* reference assemblies should note that the assembly used here, generated by Jin *et al*. ^15^, numbers chromosomal scaffolds by size (see Extended Data for synteny comparisons showing the corresponding chromosomes). Aggregating samples across the time-series, the genome-wide average signature of introgression was similar for both directions (Extended Data Fig. 5), but the chromosomal distribution of the introgression signal differed significantly between the two species (Extended Data Figs. 6,7). Local reductions in *F_ST_* and *d_XY_* in a sympatric comparison of the two species (in Brazil), relative to the allopatric comparison (non-admixed native-range populations), confirm introgression in Brazil (Extended Data Figs. 8, 9). From invasive *H. armigera* to native *H. zea*, the introgression signal is concentrated at the *CYP337B3* locus on chromosome 5, confirming previous results (Extended Data Fig. 6). Nucleotide diversity decreased over time in both donor and recipient species around *CYP337B3*, clearly demonstrating adaptive introgression (Extended Data Fig. 10). By contrast, introgression in the other direction is distributed across most chromosomes, with notable ‘plateau’ patterns consistent with locally reduced recombination on chromosomes 21, 24, and 26 (Extended Data Fig. 7). Although synteny between the two species is high, comparisons within and between both species confirm the introgression of inversion polymorphisms on these chromosomes (Extended Data Figs. 11, 12, 13, 14, 15, 16). A subset of samples sequenced using haplotagging confirm the presence of these inversions, and reveal other structural variants (SVs) that were not introgressed (Extended Data Fig. 17). Intriguingly the haploblock introgressed from *H. zea* represents the ancestral orientation in all cases (Extended Data Fig. 18). PCA-based inversion genotyping showed that all *H. armigera* individuals carrying the *H. zea* inversion allele were heterozygous (Extended Data Figs. 19, 20), suggesting a lack of strong selection. To narrow down potential instances of adaptive introgression, we next identified introgression signals that increased consistently over time.

### Temporal patterns of introgression

We divided samples into three timepoints comprising approximately equal individual sample sizes (Extended Data Fig. 21). For each we calculated *f_d_* as above, as well as the positive change in the introgression signal over the two intervals: 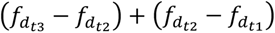. As expected, gene flow from *H. armigera* to *H. zea* increased over time at loci surrounding *CYP337B3* on chromosome 5 (Fig. 2A, Extended Data Figs. 22, 23, 24). At this locus, *H. armigera* ancestry increased rapidly among *H. zea* over the first time interval, reaching near-fixation in recently collected samples. On the remainder of chromosome 5, *H. armigera* ancestry declined over time in *H. zea* (Extended Data Fig. 24). This result is consistent with decoupling of *CYP337B3* from linked *H. armigera* alleles over successive generations, declining in frequency either through drift or selection against interspecific incompatibilities. This phenomenon is predicted in theory and can be inferred retrospectively, but is rarely observable directly.

**Figure 2.**
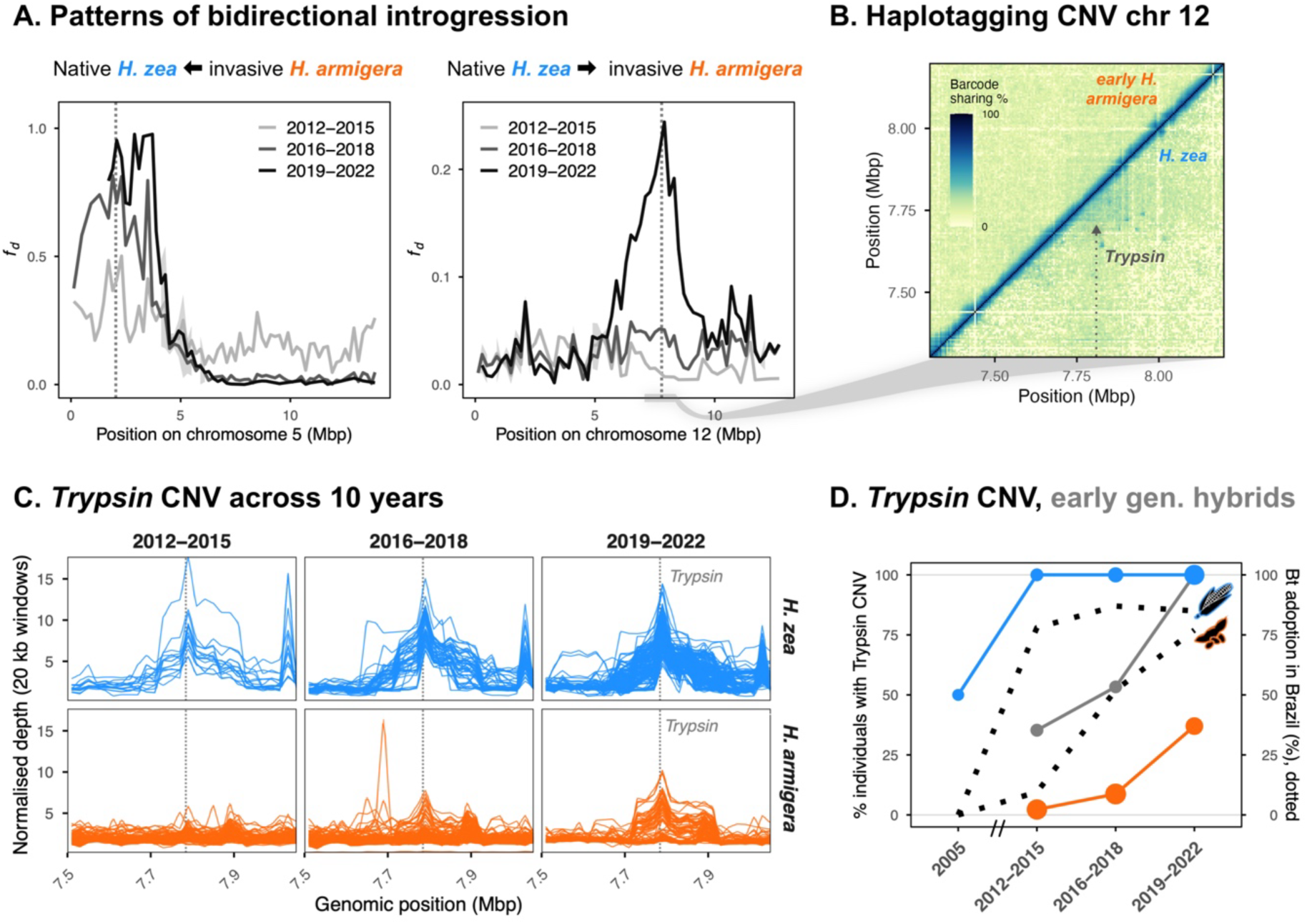
**(A)** Left panel: *f_d_* using the topology (((ZEA, ZEA_BRAZIL), ARM)), PUN), indicative of introgression from *H. armigera* to *H. zea*, shown for chromosome 5 for individuals split into three temporal sets. Right panel: *f_d_* using the topology (((ARM, ARM_BRAZIL), ZEA)), PUN), indicative of introgression from *H. zea* to *H. armigera*, shown for chromosome 12 for individuals split into three temporal sets. **(B)** Haplotag barcode sharing in *H. zea* samples collected early in the time-series (bottom right corner) and *H. armigera* collected early in the time-series (top left corner). A triangular shape consistent with a tandem duplication is evident at the introgressed trypsin locus in *H. zea* samples. **(C)** Normalised depth of coverage on chromosome 12 for *H. zea* samples (top panel) and *H. armigera* samples (bottom panel), where each individual is one line. Increases in relative depth of coverage in *H. armigera* appear over time, consistent with introgression. **(D)** Detection of the repeat cluster using depth of coverage. Circle size indicates individual sample size for *H. armigera* (orange), early-generation hybrids (grey) and *H. zea* (blue). Dashed lines: Bt adoption in Brazil; percentage of total Brazilian soy area with Bt, as reported by de Freitas Bueno *et al.* ^35^ and percentage of Brazilian maize area with Bt, as reported by Schuster *et al.* ^36^.

From native *H. zea* to invasive *H. armigera*, there was no detectable change in the frequency of *H. zea* inversion haploblocks on chromosomes 21, 24, and 26, suggesting that these structural variants are not under directional selection (Extended Data Fig. 25). Instead, introgression primarily increased over time in a narrow region of chromosome 12 in samples collected after 2018 (Fig. 2B, Extended Data Figs. 26, 27, 28). A cluster of tandem repeat trypsin genes (digestive proteases) mapped to the chromosome 12 locus showing the greatest increase in introgression from *H. zea* to *H. armigera* across the whole genome (Extended Data Fig. 27). In North American *H. zea*, this cluster has previously identified and associated with field-evolved resistance to the *Bt* toxin Cry1Ac ^28,31^. More generally, trypsins have been repeatedly implicated in Bt resistance in lepidopterans ^32–34^. We confirmed the presence of this introgressed structural variant by using a subset of linked-read haplotagged samples collected early in the time series: *H. zea* samples show a pattern of tag-sharing diagnostic of tandem duplication at the trypsin locus, while *H. armigera* samples did not (Fig. 2C). Although recently-collected *H. armigera* samples were not haplotagged, the introgressed repeat cluster could be inferred through changes in local depth of coverage over the time course (Fig. 2D). At the *Trypsin* locus, normalised depth of coverage in *H. zea* was elevated and constant between 2012 and 2022, but increased over time in *H. armigera* coincident with increased use of Bt soy in Brazil (Fig. 2D). Notably, 4 of 8 Brazilian *H. zea* samples collected in 2005 displayed the diagnostic peak in read depth, indicating polymorphism prior to the deployment of Bt in Brazil. Together these results point to selection for a structural variant conferring Bt resistance in Brazilian *H. zea* between 2005 and 2012, followed by recent introgression into *H. armigera*. We next aimed to test the hypothesis that this introgressed structural variant confers a novel resistance phenotype in Brazilian *H. armigera* exposed to Bt soy.

### A *H. zea* trypsin repeat cluster confers *Bt* resistance in wild *H. armigera*

To identify the loci underlying Bt resistance in Brazilian *H. armigera*, we collected wild individuals in 2018 and artificially selected those that survived on leaf discs of Bt soybean expressing Cry1Ac (see Methods). Tolerant and susceptible lines were crossed to produce an F2 population segregating for Cry1Ac resistance (Fig. 3A, 3B). Using whole-genome resequencing data for data for 149 F₂ individuals plus their parents and grandparents, we confirmed that all samples were *H. armigera* (minimum *H.* armigera ancestry proportion = 0.966, mean = 0.975; Extended Data Fig. 29) and carried out quantitative trait locus (QTL) mapping for Cry1Ac resistance (Extended Data Fig. 30, Extended Data Table 2). This revealed two major effect loci on chromosomes 12 and 5, which showed no evidence of epistatic interactions (Fig. 3D, Extended Data Figs. 31, 32, Extended Data Tables 2, 3, 4, 5).

**Figure 3:**
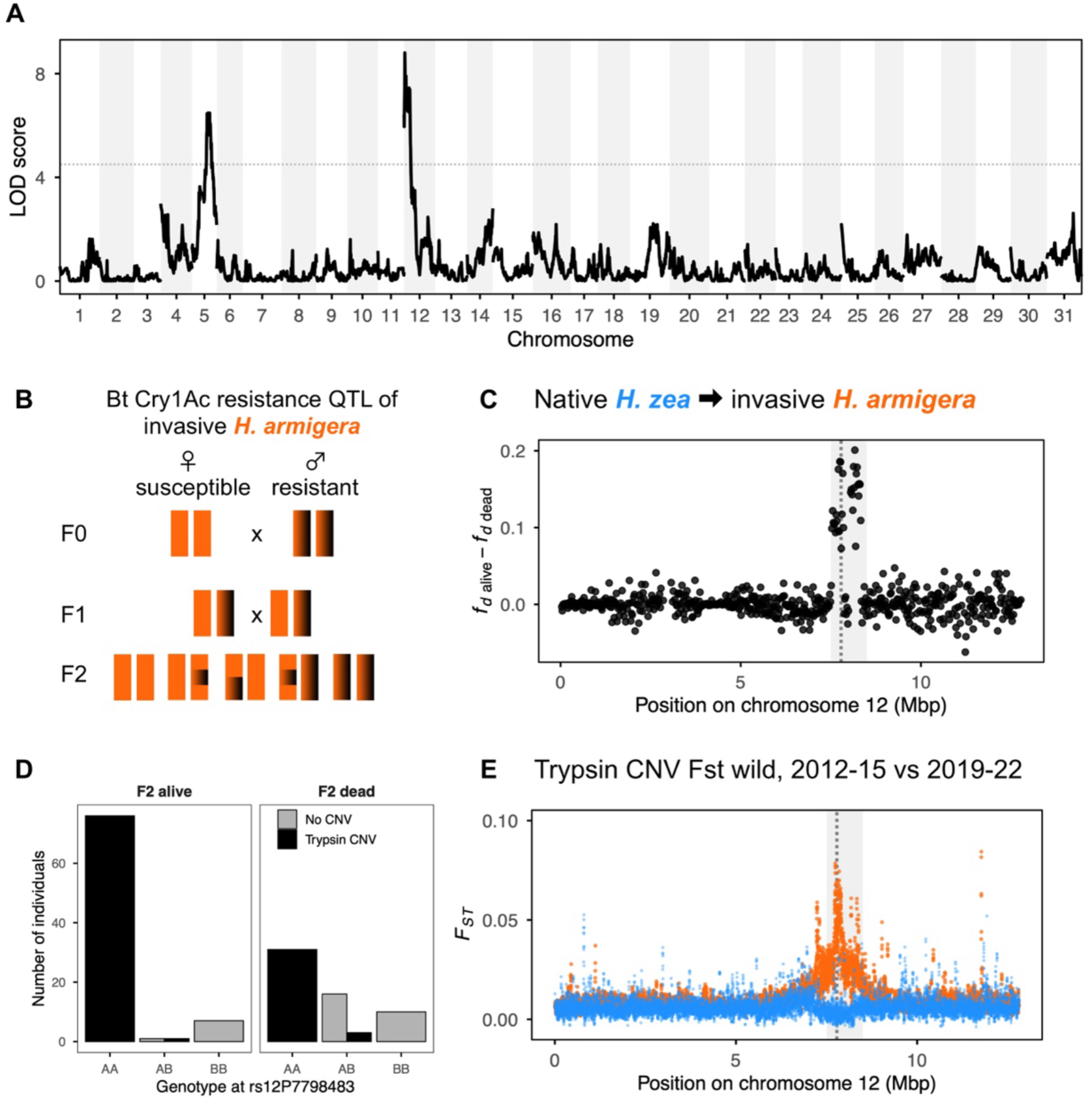
**(A)** Logarithm of odds (LOD) score for quantitative trait locus (QTL) mapping of Cry1Ac resistance in Brazilian *H. armigera*. Two major effect loci are identified on chromosomes 12 and 5 **(B)**. Intercross design for the linkage mapping experiment presented in 3A. **(C)** Difference in the signal of *H. zea* allele sharing between alive and dead individuals. In both cases *f_d_* was calculated using the topology (((ARM, QTL_SAMPLE), ZEA)), PUN). Dashed line indicates the trypsin locus, with grey shaded regions indicating the QTL confidence intervals. *H. armigera* samples that survived when exposed to Cry1Ac carried an excess of *H. zea* alleles on a haplotype directly overlapping with the QTL confidence interval. **(D)** Count of individuals in the intercross that survived or died when exposed to Cry1Ac, split by genotype and detection of the trypsin CNV based on excess normalised depth of coverage. The A allele, associated with survival in homozygotes, corresponds to the trypsin CNV. **(E)** Genetic differentiation (*F_ST_* calculated using genotype likelihoods) on chromosome 12 between samples collected early and late in the time-series. *H. armigera* is shown in orange; *H. zea* is shown in blue. *H. armigera* samples show excess differentiation at the trypsin locus (dashed line; QTL confidence intervals shaded in grey), consistent with introgression over time. *H. zea* shows reduced differentiation at this locus compared to the rest of the chromosome, consistent with a selective sweep.

We next quantified introgression signals as above for the *H. armigera* QTL samples. Remarkably, both major-effect loci showed a localised enrichment of *H. zea* ancestry among individuals that survived Cry1Ac treatment (Fig. 3C, Extended Data Fig. 33). This confirms predictions that locally adapted *H. zea* variants would enhance *H. armigera* fitness in Brazil ^20,23^. However, only one of the two QTL showed evidence of introgression in field-collected samples. At the chromosome 5 QTL (LOD= 6.48, *p*=3.29×10^−7^) there were no obvious signs of interspecific introgression in wild-caught *H. armigera* (Extended Data Fig. 33) despite its additive effect on Cry1Ac resistance. In contrast, confidence intervals for the chromosome 12 QTL (LOD= 8.80, *p*=1.57×10^−9^) overlapped with the *Trypsin* repeat cluster (Fig. 3C), which we identified as the locus with the strongest increase in introgression through time in the wild (Fig. 2A). *Trypsin* CNV presence/absence genotyping based on depth of coverage at this locus showed that individuals with the CNV were significantly more likely to be Bt-resistant (Fig. 3D). In wild *H. armigera*, this same locus showed extreme genetic differentiation over time relative to the genome-wide average (Fig. 3E, Extended Data Fig. 10). Therefore, laboratory selection for Cry1Ac resistance in Brazilian *H. armigera* had the effect of enriching for existing introgressed *H. zea* structural variant alleles, thereby replicating the adaptive introgression observed in wild populations. Interestingly, this QTL showed signs of underdominance in the presence of Bt, where fitness is lowest in heterozygotes (Extended Data Fig. 32). Despite this apparent barrier to introgression at low frequencies, the introgressed trypsin allele frequency in *H. armigera* closely tracks the rollout of Bt soy in Brazil ^35^ (Fig. 3D). Although the CNV was fixed in *H. zea* collected from 2012 onwards, samples from 2005 were polymorphic. This suggest that *H. zea* evolved resistance relatively recently, likely through gene flow from North American *H. zea* populations exposed to Cry-expressing maize and cotton since the late 1990s, followed by adoption in Brazilian maize since 2007 (Fig. 3D).

## Discussion

Together these findings indicate persistent but permeable species boundaries upon secondary contact between native and invasive agricultural pests, resulting in bidirectional adaptive introgression of structural variants conferring resistance. In addition to adaptive introgression of *CYP337B3* from invasive *H. armigera* to native *H. zea*, more recent adaptive introgression in the other direction has introduced Bt (Cry1Ac) resistance alleles to ~30% of the invasive *H. armigera* population. Thus, hybridization has generated a novel combination of traits — soy host plant adaptation in *H. armigera* and Bt resistance selected in *H. zea* — which together enable colonisation of an expansive anthropogenic habitat: Brazilian Bt soybean monoculture.

Just as ancient hybridization can combine old adaptive variants to facilitate adaptive diversification over thousands of years, these results demonstrate how contemporary hybridization can assemble independently selected alleles into a single admixed genome, resulting in adaptation over ecological timescales. Notably, the recombined adaptive variants do not need to be ‘ancient’: the trypsin allele underlying Cry1Ac resistance was polymorphic in Brazilian *H. zea* as recently as 2005, and has only recently been selected — likely through exposure to Cry-expressing maize and cotton in North America ^28^. Thus, anthropogenic change not only drives rapid adaptation through selection on standing variation, but can also promote hybridization, which can further accelerate adaptation. Consequently, in this system, resistance to control measures has evolved in two major agricultural pests through bidirectional adaptive introgression over the course of less than a decade.

The practical outcome of hybridization between *H. armigera* and *H. zea* is the ‘worst of both worlds’: two distinct agricultural pests are now resistant to control measures, and show greater adaptive potential. In the case of *H. zea*, the pyrethroid resistance gene *CYP337B3* had reached near-fixation in the most recently collected samples. This large pyrethroid-resistant Brazilian source population of *H. zea* is a biosecurity threat to the two largest maize producing agro-economies: the United States, where the spread *H. armigera*-derived *CYP337B3* in *H. zea* is already causing an increase in pyrethroid control failures ^37^, and China, where *H. zea* is non-native. In the case of *H. armigera*, approximately 30% of samples collected between 2019 and 2022 carried Cry1Ac resistance alleles. An additional *H. zea*-derived Cry1Ac resistance QTL on chromosome 5 was present in wild-caught *H. armigera*, and is expected to spread in the population given its additive phenotypic effect. These results provide an early warning of Cry1Ac-resistant *H. armigera* in Brazil. More broadly, Brazilian *H. armigera* pose an increasingly imminent biosecurity threat to the U.S., where the species has not yet spread. Transport of Brazilian *H. armigera* back into its native range is also a biosecurity concern. For example, Australian *H. armigera* populations harbour monogenic resistance mutations for the Bt toxin Vip3Aa ^38^. This raises the possibility of recombinant trans-Pacific *H. armigera* genotypes conferring resistance to multiple Bt toxins, potentially threatening ‘pyramiding’ strategy of multi-toxin Bt crop use.

Overall, a combination of serial-sampled genomic data and linkage mapping has revealed a combinatorial mechanism driving rapid adaptation. The outcome of interspecific hybridization can range from rapid adaptation to demographic collapse, ultimately depending on the genetic architecture and ecological contingency of hybrid fitness ^39^. In systems such as this, admixture has enabled the sharing of adaptive variation between distinct and competing species. For pests and pathogens, such rapid evolutionary rescue presents a challenge to food security and multidrug resistance. For some threatened taxa, the same process could facilitate persistence. Recognising hybridization as a driver for adaptation on anthropogenic timescales will be essential for understanding evolutionary responses to global change.

## Methods

### Sample collection, DNA extraction, library preparation, and sequencing

For samples in ‘batch 1’, *Helicoverpa* adults were collected throughout Brazil as part of routine monitoring. For all other sample batches, *Helicoverpa* larvae were collected from agricultural regions across Brazil and sent to the Arthropod Resistance Laboratory at the Luiz de Queiroz College of Agriculture, University of São Paulo, as part of routine monitoring of susceptibility to chemical insecticides and Bt proteins during the 2012–2022 crop seasons (Fig. 1A, Extended Data Table 1; SISBIO numbers: 65052, 75582 and 61824; SisGen numbers: A792B04 and R1EA05A; TTM/ATM number: 1014273). For each field-collected sample, we performed both molecular identification to distinguish *H. armigera* from the native *H. zea*, following the PCR–RFLP protocol described by Behere *et al.* ^1^. Field-collected larvae were reared for at least one generation under controlled laboratory conditions (25 ± 1°C, 70 ± 10% RH, and a 14h photophase). Between three and seven days after adult emergence, individuals from each population in each season were individually placed in 1.5mL Eppendorf tubes containing 98% ethanol and stored at –20 °C.

Genomic DNA was extracted from 2mm wide segments of the thorax using Kučka and Chan’s high molecular weight magnetic bead DNA extraction protocol ^2^. DNA concentration was quantified using a Qubit™ fluorometer (Invitrogen, USA), and low-concentration samples were re-extracted or excluded. For a subset of samples, DNA was extracted either from the thorax or leg using the CTAB extraction protocol and resuspended in 50 µl of Milli-Q water ^3^. For this subset, DNA quality control was initially assessed by visual inspection on a 0.8% (w/v) agarose gel, followed by spectrophotometric and fluorometric analyses, and each sample was evaluated using a NanoDrop spectrophotometer to detect potential contaminants and an agarose gel to assess DNA integrity, and subsequently quantified using a Qubit fluorometer.

For all samples, sequencing libraries were prepared via a tagmentation based approach using a homemade tn5 transposase as per Picelli *et al.* ^4^. Prior to pooling, DNA concentration and insert size were assessed using a Qubit™ fluorometer and TapeStation (Agilent Technologies) respectively. Equal amounts of each library were pooled together, and then sequenced (150bp paired-end) using an Illumina NovaSeq 6000 (Novogene UK).

#### Haplotagging Library Construction and Sequencing

Haplotagging libraries were prepared following the protocol of Meier *et al.* ^5^. In total, we analysed 93 individuals: 63 *H. armigera*, 16 *H. zea*, and 14 putative hybrids collected in Brazil between 2012–2018. Genomic DNA was extracted from thorax tissue and normalised to 0.15 ng µl^−1^ in 10 mM Tris (pH 8). Libraries were constructed in 96-well plates. Each sample received 1.2 µl of haplotagging beads (~0.9 million beads carrying one of 885K barcodes), 30 µl WASH buffer (20 mM Tris pH 8, 50 mM NaCl, 0.1 % Triton-X100), 10 µl of 5× tagmentation buffer (50 mM TAPS pH 8.5, 25 mM MgCl₂, 50 % DMF), and 25 µl of 0.6 % SDS to strip Tn5 after tagmentation.

For amplification, one-tenth of the bead–DNA complexes from each of the 96 samples was pooled within eight-tube strips, and subsequently collapsed into four final pools. After placing the bead pools on a magnetic rack, residual buffer was removed and replaced with 20 µl of 1× Lambda Exonuclease buffer supplemented with 10 U exonuclease I and 5 U Lambda exonuclease (NEB). Beads were incubated at 37 °C for 30 min, washed twice with 150 µl WASH buffer, and resuspended for PCR. Libraries were amplified using Q5 High-Fidelity DNA polymerase (NEB) in four 25 µl reactions per pool, using 10 µM TruSeq-F (AATGATACGGCGACCACCGAGATCTACAC) and TruSeq-R primers (CAAGCAGAAGACGGCATACGAGAT). Cycling conditions were: 72 °C for 10 min; 98 °C for 30 s; followed by 10 cycles of 98 °C for 15 s, 65 °C for 30 s, and 72 °C for 60 s. PCR products were pooled, size-selected using AMPure XP beads, quantified by Qubit, and normalised to 2.5 nM using 10 mM Tris (pH 8.0, 0.1 mM EDTA).

Pooled libraries were sequenced on an Illumina HiSeq 3000, using a 150 + 13 + 12 + 150 (R1 + i7 + i5 + R2) configuration. This design produces 13-bp i7 and 12-bp i5 index reads corresponding to haplotag plate and well barcodes.

### Bioinformatics

#### Read mapping

The mapping and variant-calling pipeline was executed using Snakemake ^6^. We mapped reads to the highest-quality available *H. armigera* reference assembly, generated from a sample collected in north-western China ^7^. After trimming fastq files using Fastp v0.23 ^8^, and bwa mem v0.7.12 ^9^ was used to map reads. SAMtools v1.10 ^10^ was used to sort and index the resulting bam files. Duplicates were marked and removed using Picard’s MarkDuplicates v2.9.2 ^11^.

#### Sequencing Depth Profiling and Sexing

To characterise genome-wide and chromosome-specific sequencing depth per individual, we used mosdepth v0.3.3 and d4tools v0.3.4 to compute and summarise base-level coverage from mapped reads. All analyses were conducted on BAM files previously aligned to the reference genome. Depth was computed per base using mosdepth with the following parameters: --mapq 20 to restrict to confidently mapped reads, --d4 to output in compressed D4 format, and --fast-mode to skip insertions and deletions. Each sample was processed in parallel using 4 threads. To obtain per-chromosome depth summaries, we ran d4tools stat with the -s median and -s mean options on the .d4 files. This produced one summary file per sample containing median and mean depth values per chromosome. These values were used in downstream analyses to infer sex chromosome copy number, normalise depth across the genome, and examine population-level variation in coverage patterns.

#### Haplotagging Read Processing, Mapping and SV detection

Raw BCLs were converted to FASTQ using bcl2fastq v2.20 with: --use-bases-mask Y150,I13,I12,Y150 --minimum-trimmed-read-length 1 --mask-short-adapter-reads 1 --create-fastq-for-index-reads (Illumina). Haplotag barcodes consist of four 6-bp sequences embedded in the FASTQ read name. Barcode strings were assigned to the BX tag following Meier *et al.* ^5^, allowing one mismatch when an unambiguous closest match existed. Adapter trimming and quality filtering (if required) were conducted prior to alignment. Reads were mapped to the H. armigera reference genome (GenBank: GCA_030705265.1) using BWA-MEM2, retaining barcode information using the –C flag. Alignments were coordinate-sorted with samtools, and PCR duplicates were marked with Picard MarkDuplicates using: READ_ONE_BARCODE_TAG=BX CREATE_INDEX=true. This preserves molecule-level identity during duplicate marking. All resulting BAMs were inspected to confirm correct propagation of BX:Z: tags and uniform coverage across windows. These processed, barcode-aware BAM files were used for downstream analyses.

We identified candidate structural variants (SVs) using Wrath v2024.03, a tool designed to detect large-scale SVs from linked-read sequencing data by quantifying barcode sharing between genomic windows ^12^. Analyses were performed separately for each chromosome using the BAM files with barcodes encoded in the BX tag. To run Wrath, we provided a list of BAM files per group, the chromosome of interest and a fixed window size of 10 kb. Wrath calculates the Jaccard index of shared barcodes between each pair of non-overlapping windows, producing a barcode-sharing matrix for each chromosome.

#### Variant calling for call sets 1 and 2

In addition to genotype-liklihood-based methods, some analyses made use of variant calls. Variants were called using GATK HaplotypeCaller v4.3 ^13^. To improve genotyping calling performance, and to enable comparison with reference populations, a diverse array of publicly available *Helicoevrpa* sp. was used, in addition to samples from closely related species. Before filtering, a total of 1366 individuals were included for genotyping. These included 1017 newly sequenced Brazilian *Helicoverpa* samples; 124 native-range *H. armigera* reported by Jin *et al.* ^7^, of which 20 were used as reference non-admixed *H. armigera*; a further 31 *Helicoverpa* samples reported by Anderson *et al.* ^14^ including 7 *H. punctigera* individuals as an outgroup; 13 *H. zea* sampled in 2002 reported by Taylor *et al.* ^15^ for use as reference non-admixed samples; 4 *H. zea* samples collected in the US in 2019 by North *et al.* ^16^; 62 *Helicoverpa* samples collected in North America by North *et al.* (2026); 112 Brazilian *Helicoverpa* samples collected by Valencia Montoya *et al.* ^17^ and Anderson *et al.* ^14^; and three additional outgroups: *Chloridea subflexa* and *C. virescens* generated by Guo *et al.* ^18^ and *H. assulta* generated by Xu *et al.* ^19^.

Some newly sequenced samples previously sequenced by Valencia Montoya *et al.* ^17^ or Anderson *et al.* ^14^. Of these duplicates, we retained whichever had the highest depth. After removing such duplicates, we retained a total of included 1128 Brazilian *Helicoverpa* samples, including those used for linkage mapping.

GATK CombineVCFs was used to merge GVCF files across individuals and GenotypeGVCFs was used to call genotypes jointly across samples. In order for genotyping to be computationally feasible, each of the 31 chromosomes were divided into 20 equally sized intervals. Genotyping was carried out in parallel per-interval before concatenating intervals into a single VCF file.

Two filtering approaches were used to generate call set 1 and call set 2, respectively.

#### Filtering: call set 1

Call set 1 was used to quantify introgression, and was applied to Brazilian *Helicoverpa* samples, and reference non-admixed groups (see Quantifying Introgression, below). First, bcftools v1.19 was used to retain only biallelilc SNPs from Brazilian *Helicoverpa* samples with an autosomal mean depth of coverage of at least 5X, as calculated using vcftools v 0.1.17. Reference non-admixed samples were also required to meet this criteria, with the exception of 5 *H. punctigera* samples (M0236, M0239, M0244. M0245, and M0263) which fell marginally below an average of 5X. From here, all samples GATK VariantFiltration and SelectVariants (v4.2.5.0) were used to flag and exclude variants according to the following conditions: SOR > 3.0, FS > 60.0, MQRankSum < −20.0, MQRankSum > 20.0, ReadPosRankSum < −10.0, ReadPosRankSum > 10.0. Invariant sites were then excluded. Next, vcftools v 0.1.17 was used to apply the following conditions: --max-missing 0.75 --minQ 20 --min-meanDP 5 --minDP 5. Next, individual-level genotype-level filters were applied using a custom python script (filter1.py), which retained genotypes with a depth in the range between 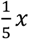 and 5*x* where *x* is the mean autosomal depth of coverage for that individual calculated using vcftools, and otherwise set genotype calls to missing data (‘./.’). Finally, after the introduction of missing data through the genotype-level filter, vcftools --max-missing 0.75 was re-applied. This resulted in a VCF file of 628 individuals containing 25,319,153 sites across all chromosomes.

#### Filtering: call set 2

Call set 2 consisted of more individuals and a less-stringent filter with respect to the proportion of missing data and the per-individual mean depth of coverage. Call set 2 was only used for individual-level summary statistics (see Ancestry Proportion and Ancestry Heterozygosity, below). Filter 2 was only to autosomal loci for Brazilian *Helicoverpa* samples and reference non-admixed samples with a depth of at least 0.5X using mosdepth (see above). First, GATK VariantFiltration and SelectVariants was used to apply the following conditions: SOR > 3.0, FS > 60, MQRankSum < −20.0, MQRankSum > 20.0, ReadPosRankSum < −10.0, ReadPosRankSum > 10.0. Next, vcftools was used to apply the following filters: -max-missing 0.5 --minQ 20 --min-meanDP 5 --minDP 5. Next, individual-level genotype filters were applied using a custom python script (filter2_4_v2.py) using the same logic as that used for filter 1: retaining genotypes with a depth in the range between 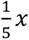 and 5*x* where *x* is the mean autosomal depth of coverage for that individual. Finally, vcftools --max-missing 0.5 was re-applied. This resulted in a VCF file of 1,014 individuals comprising 31,013,075 polymorphic autosomal loci.

#### Ancestry proportion ancestry heterozygosity using call set 2

The ancestry proportion (AP, equivalent to the hybrid index) and ancestry heterozygosity (AH, equivalent to the interclass heterozygosity) were calculated using sites that were fixed differences when comparing reference non-admixed *H. armigera* and reference non-admixed *H. zea* by applying scripts adapted from Liu *et al.* ^20^ to call set 2. Specifically, the AP is the per-sample proportion of alleles derived from *H. armigera*, while the AH is the proportion of genotypes that are heterozygous. AH is therefore equivalent to Fitzpatrick’s interclass heterozygosity *H_I_* and Turelli and Orr’s *p*), while the AP is directly comparable to the hybrid index calculated in genomic or geographic cline analyses ^21,22^. The AP correlated strongly with the ANGSD ancestry proportion (Extended Data Fig. 1).

#### Quantifying introgression using call set 1

Given the availability of samples collected in locations and times when admixture had never been reported (native-range *H. armigera* collected in North-Western China and *H. zea* collected in North America in 2002, and *H. punctigera*), the following ‘backbone’ phylogeny could be used: (ARM, ZEA), PUN. Using the ANGSD *H. armigera* ancestry proportion to assign Brazilian *Helicoverpa* samples to *H. armigera* (>0.5) or *H. zea* (<0.5), introgression could therefore be measured in both directions. We reasoned that gene flow from invasive *H. armigera* to native Brazilian *H. zea* would result in an excess of ABBA patterns relative to BABA in the following topology: (((ZEA, ZEA_BRAZIL), ARM)), PUN). In the reverse direction, gene flow from native *H. zea* to invasive Brazilian *H. armigera* would result in an excess of ABBA patterns relative to BABA in the following topology: (((ARM, ARM_BRAZIL), ZEA)), PUN). Using this framework, the summary statistic *f_d_* ^23^ were calculated in 200kbp windows with at least 500 informative sites, and 20kbp windows with at least 200 usable sites. This was done for both test topologies using Python scripts described in Martin *et al.* ^23^. For window-based analyses, outlier regions were defined as the upper 5^th^ and 1^st^ percentile of the *f_d_* distribution.

Samples were divided into three timepoints comprising approximately equal individual sample sizes (Extended Data Fig. 21). For each, the positive change in the introgression signal over the two intervals was calculated as 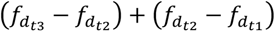.

Additionally, *F_ST_* and *d_XY_* were calculated for the same window sizes using the same informative site threshold for two comparisons: an allopatric comparison (reference non-admixed *H. armigera* and *H. zea*) and a sympatric comparison (Brazilian *H. armigera* and *H. zea*). This was used to confirm the expectation that regions introgressed in sympatry should show high *f_d_* values and reduced *F_ST_* and *d_XY_* in sympatry relative to allopatry.

#### Mapping and genotyping inversion polymorphisms using call set 1

DGENIES, which implements Minimap2, was used to generate genome-wide dot plot synteny maps and MAF alignment files ^24,25^. Only query and reference alignments of at least 200kbp, and assembly scaffolds of at least 1Mbp, were retained. Synteny plots (Extended Data Figs. 11, 12, 13, 14, 15, 16) were generating using R scripts described by Kim *et al.* ^26^. This was applied to intra- and inter-specific comparisons between *H. armigera* and *H. zea* reference assemblies ^7,27–30^.

To assign inversion genotypes, PCA was calculated with PLINK using only loci within the inversion regions of interest, as defined by the breakpoints mapped with Minimapper2. Clusters were assigned to genotypes manually according to expected genotypic states under the model proposed in Extended Data Fig. 18.

#### Population Structure Inference from Genotype Likelihoods

Population structure was inferred using genotype likelihoods computed in ANGSD v0.935, followed by principal component analysis and admixture estimation using PCAngsd v1.10. BAM files were grouped by population subset and filtered to retain sites where ≥80% of individuals were genotyped. Genotype likelihoods were computed using the GATK model (-GL 2) with base and mapping quality filters (-minQ 20, -minMapQ 30), and with major alleles and allele frequencies estimated internally (-doMajorMinor 1, - doMaf 1). Depth filters were scaled by the number of individuals: a minimum of 2×N and a maximum of 5×mean depth×N. Genotype likelihoods were output in Beagle format (-doGlf 2) and analysed with PCAngsd using the following non-default parameters: number of ancestral populations fixed at K=2, shrinkage of the covariance matrix enabled, and no LD pruning. PCAngsd was run with --admix, --beagle, and inferred admixture proportions and principal components directly from the genotype likelihoods.

#### Admixture proportions using genotype likelihoods

Population structure and ancestry proportions were inferred using PCAngsd v1.10, which estimates covariance matrices and admixture components directly from genotype likelihoods. Genotype likelihoods were computed in ANGSD using the filtering scheme described above, but without downsampling sites; instead, we retained all genomic positions that passed quality filters and for which ≥75% of individuals in the population had at least one informative read (ANGSD option -minInd, set to 0.75 × *N*). This approach preserves the full genomic signal while excluding sparse or poorly covered sites that could bias covariance estimation. For each population, genotype likelihoods were exported in Beagle format after applying minimum allele frequency (-minMaf 0.05), site-quality, and depth filters, as well as a population-specific maximum-depth threshold to avoid inflated likelihoods from read pile-ups. Concatenated per-chromosome likelihood files were used as input to PCAngsd, which jointly estimates the covariance matrix under a probabilistic framework and computes individual ancestry coefficients without the need for called genotypes. The resulting admixture coefficients were taken from the .*admix* output and used as continuous ancestry estimates for all downstream analyses, including Fst contrasts and temporal structuring.

#### Detection of hybridisation hotspots across space and time

To test whether hybrids overlap in space and time with regions where both species (*H. armigera* and *H. zea*) were sampled at a given timepoint, we clustered individuals by geographic proximity within each year. Specifically, we applied hierarchical clustering using average linkage on geographic coordinates (latitude and longitude), grouping individuals sampled within the same year into spatial clusters with a maximum pairwise distance of 100 km (using the Haversine distance). This approach accounted for the high dispersal ability of Helicoverpa moths while ensuring sufficient sample sizes within clusters. For each year-specific cluster, we recorded the number of individuals assigned to each of three ancestry categories: H. armigera (>95% inferred ancestry), H. zea (<5%), and early-generation hybrids (intermediate ancestry). A cluster was classified as an area of species overlap if it contained at least three individuals of each parental species (H. armigera and H. zea). We then quantified the number of clusters with and without overlap, and with and without hybrids, and visualised their distributions as a stacked barplot. To test whether overlap between the parental species was associated with the presence of hybrids, we performed a Chi-squared test of independence with Yates’ continuity correction.

#### Genetic Differentiation Analyses using genotype likelihoods

Genetic differentiation between population pairs was estimated using the site allele frequency (SAF)–based FST framework implemented in ANGSD v0.935 and realSFS. Individuals were grouped into predefined groups by time ranges (12-15, 16-18, 19-22) and species’ ancestries (considered one of the two study species in their ancestry proportion was >95%). Genotype likelihoods were computed using the SAMtools model (-gl 1) with filters for read and mapping quality (-minQ 20, -minMapQ 30), and base alignment quality recalculation enabled (-baq 2). Sites were further filtered to remove low-quality reads (-remove_bads 1) and counts were generated using -doCounts 1. SAFs were estimated per population using -dosaf 1, with ancestral and reference sequences both specified (-anc, -ref). To avoid bias from over-represented sites, a maximum depth threshold scaled by population size and average depth was applied (-setMaxDepth), calculated as 5×mean depth×N. A joint two-dimensional site frequency spectrum (2D-SFS) was estimated between population pairs using realSFS, based on their .saf.idx likelihood files. This 2D-SFS was then used to compute FST indices with realSFS fst index, and genome-wide sliding window FST values were calculated with realSFS fst stats2 using a window size of 5 kb and step size of 1 kb for plotting.

#### Nucleotide Diversity and Neutrality Statistics using genotype likelihoods

Nucleotide diversity (*θ* statistics) was estimated for each predefined population using the ANGSD–realSFS workflow. For each population, genotype likelihoods were recalculated as described above and per-population site allele frequency (SAF) likelihoods were generated using -dosaf 1, with the reference genome also specified as the ancestral state. To avoid inflation of diversity estimates due to read pile-ups, a population-specific maximum depth threshold was applied using -setMaxDepth, calculated as 5 × mean depth × number of individuals in the population. For each chromosome (1–31), an unfolded per-chromosome site frequency spectrum (SFS) was first inferred with *realSFS*, restricted to the corresponding genomic interval. This SFS was then used to convert SAF likelihoods into *θ*-estimators with *realSFS saf2theta*, generating per-site estimates of pairwise nucleotide and Tajima’s *D* among others. Genome-wide summary statistics for each chromosome were obtained using thetaStat do_stat. Sliding-window diversity profiles were then computed for each chromosome with thetaStat do_stat -win 50000 -step 10000, producing 50 kb windows stepped every 10 kb.

### Linkage mapping

#### Phenotype bioassay

A total of 395 F₂ intercross larvae were tested on fresh Bt soybean leaves expressing the insecticidal protein Cry1Ac. Of these, 303 larvae survived (77%) and 92 larvae died (23%). Following DNA extraction and library preparation, due to the low quality of some DNA samples, we successfully sequenced 87 surviving larvae (58%) and 62 dead larvae (42%), resulting in a total of 149 sequenced larvae from the F₂ intercross population.

#### Linkage map

The linkage map was constructed using Lep-MAP3 ^31^, utilizing genotypic data from 149 F₂ individuals along with their parents and grandparents, totalling 3,738 markers. The “ParentCall2” module (with the option removeNonInformative=1) was employed to estimate the most accurate posterior probabilities of parental genotypes. Additionally, we used the “Filtering2” module (with the option dataTolerance=0.01) to remove markers exhibiting significant segregation distortion. The modules “SeparateChromosomes2” and “joinSingles2all” were not utilized because the reference genome is assembled at the chromosome level; thus, markers were retained on their respective chromosomes.

We ran the “OrderMarker2” module for each of the 31 chromosomes 15 times and selected the run with the highest likelihood score. In this step, specific options such as “outputPhasedData=1”, “recombination2=0”, “usePhysical=1 0.1” and “useKosambi=1” were applied. The “OrderMarker2” module orders markers within each linkage group by maximizing the likelihood of the data given the marker order. Markers that did not show strong linkage with their flanking markers were manually removed to avoid long gaps. The final linkage map provided phased chromosomal marker data with missing genotypes imputed. The phased data were converted for QTL mapping using Lep-MAP3’s map2genotypes.awk script, and the genotypes 1 1, 1 2, 2 1, and 2 2 were transformed to AA, AB, AB, and BB, respectively.

The final linkage map consisted of 3,735 markers distributed across 31 linkage groups (LGs), covering a total length of 3,661.8 centimorgans (cM). The number of markers per LG ranged from 95 to 156, and LG lengths varied between 88.3 and 143.6 cM. The average inter-marker spacing within each LG ranged from 1.4 to 14.2 cM (Extended Data Fig. 30 and Table S2).

#### QTL Mapping

QTL mapping was performed using the R/qtl package ^32^ in the R program ^33^, employing the Expectation-Maximization (EM) algorithm implemented in the “scanone” function. The phenotypes “survival” and “death” were coded as “1” and “0”, respectively, and the parameter model = “binary” was specified to accommodate the binary nature of the trait. The functions “fitqtl” (with drop-one-term analysis) and, subsequently, “refineqtl” were used to identify QTL positions with maximum likelihood under a fixed QTL model. The significant LOD threshold (0.95 percentile) was determined through 10,000 permutation tests. Confidence intervals for the detected QTLs were estimated using the “lodint” function. Additionally, we used the “scantwo” function to perform a two-dimensional genome scan with a two-QTL model to detect potential epistatic interactions between QTLs.

The QTL analysis for the survival trait revealed suggestive peaks (LOD > 4.09) (Extended Data Table 3 and Fig. 1). Two significant QTLs associated with survival were identified, one on LG 5 (LOD = 6.48, *P* < 0.05) and another on LG 12 (LOD = 8.80, *P* = 0.004) (Extended Data Fig. 31). Additionally, a peak was observed on LG 12 at position 104.9 cM. However, the QTL analysis did not support this peak (LOD = 0.51, *P* = 0.303). The QTL on LG 5 exhibited a confidence interval of 15.18 cM, while the QTL on LG 12 had a confidence interval of 9.65 cM (Extended Data Table 3).

An epistatic interaction test between the QTLs on LGs 5 and 12 showed no significant interaction (LOD = 1.42, *P* = 1), indicating that these QTLs do not exhibit epistatic effects associated with the survival trait (Extended Data Table 4). Individuals carrying the heterozygous genotype of the QTL on LG 5 displayed intermediate survival, indicative of incomplete dominance. In contrast, the survival was lowest among those with the heterozygous genotype of the QTL on LG 12, suggesting underdominance (Extended Data Figure 32 and Extended Data Table 5).

#### Identification of Candidate genes

To investigate potential genes associated with *H. armigera* survival in soybeans that express Cry1Ac protein, we used a 300 kb upstream and downstream of regions near each significant QTL marker identified. This process utilized the genome annotation file (GCF_030705265.1-RS_2024_03) from *H. armigera* genome assembly GCF_030705265.1 published in the NCBI.

## Data Availability

All newly generated sequencing data have been uploaded to the ENA: ERP162059. Sample metadata is available in Table S1 and on the Earthcape database: https://heliconius.ecdb.io/

## Code Availability

Custom scripts available at https://github.com/Insect-evolution-and-genetics-lab/genome_collision_scripts

## Acknowledgements

We are grateful to colleagues for their comments and feedback, especially Joana Meier and members of the Rapid Speciation Group at the Sanger Institute, and all members of the Insect Evolution and Genomics Group at Cambridge between 2019 and 2025.

Jochen Wolf and Andrea Manica also provided thoughtful comments on preliminary results. HN was funded by the JS Gardiner Fund as a PhD student.

## Author contributions

**HLN**: Conceptualisation (computational analysis, sampling design), data curation, formal analysis (bioinformatics), methodology (bioinformatics), project administration, software implementation, investigation (lab work). **GMK**: Conceptualisation (computational analysis, sampling design), data curation, formal analysis (bioinformatics), methodology (bioinformatics), project administration, software implementation, supervision (of AW), investigation (lab work). **DA**: Conceptualisation (computational analysis, experimental design), data curation, formal analysis (bioinformatics, linkage map), methodology development and implementation (DNA extraction and library preparation), project administration, investigation (bioassay), software implementation (QTL mapping), resources (sample provision). **IAW:** Methodology (DNA extraction and library preparation), investigation (lab work), data curation. **MK:** Methodology (DNA extraction, library preparation, haplotagging), investigation (lab work). **AW:** formal analysis (spatial analysis). **YFC:** Conceptualisation, Resources, Funding. **TW:** Conceptualisation, Resources, Funding, Data Curation. **ASC:** Conceptualisation, Investigation (DNA extraction and Library preparation), Formal Analysis (preliminary analysis), Resources, Funding, Data Curation, Project Administration. **CO:** Conceptualisation, Resources, Funding, Data Curation, Project Administration. **CDJ:** Conceptualisation, Supervision (of HLN, GMK, DA, AW), Resources, Funding, Data Curation, Project Administration.

## Extended Data

Citations used in Extended Data Tables and Figures correspond to those in the Methods References.

**Extended Data Table 2.** Linkage map summary.

| LG | Nº Marker | Length | Ave. Spacing | Max. Spacing |
| --- | --- | --- | --- | --- |
| 1 | 119 | 141.5 | 1.2 | 6.9 |
| 2 | 127 | 122.8 | 1.0 | 4.8 |
| 3 | 133 | 96.6 | 0.7 | 2.1 |
| 4 | 119 | 112.4 | 1.0 | 3.5 |
| 5 | 124 | 88.3 | 0.7 | 1.4 |
| 6 | 127 | 93.8 | 0.7 | 1.4 |
| 7 | 141 | 138.0 | 1.0 | 6.9 |
| 8 | 136 | 125.6 | 0.9 | 8.4 |
| 9 | 119 | 109.7 | 0.9 | 4.1 |
| 10 | 128 | 110.4 | 0.9 | 3.5 |
| 11 | 112 | 93.8 | 0.8 | 4.1 |
| 12 | 130 | 111.8 | 0.9 | 3.5 |
| 13 | 122 | 114.5 | 0.9 | 3.5 |
| 14 | 117 | 91.7 | 0.8 | 3.5 |
| 15 | 130 | 143.6 | 1.1 | 6.2 |
| 16 | 125 | 135.3 | 1.1 | 5.5 |
| 17 | 116 | 97.3 | 0.8 | 2.8 |
| 18 | 117 | 115.9 | 1.0 | 4.8 |
| 19 | 156 | 141.4 | 0.9 | 2.8 |
| 20 | 137 | 142.8 | 1.1 | 3.5 |
| 21 | 121 | 127.0 | 1.1 | 5.5 |
| 22 | 121 | 111.1 | 0.9 | 3.5 |
| 23 | 110 | 95.9 | 0.9 | 3.5 |
| 24 | 110 | 137.4 | 1.3 | 5.5 |
| 25 | 101 | 120.8 | 1.2 | 6.2 |
| 26 | 112 | 104.9 | 0.9 | 4.8 |
| 27 | 98 | 133.3 | 1.4 | 5.5 |
| 28 | 119 | 125.6 | 1.1 | 4.8 |
| 29 | 100 | 123.6 | 1.2 | 6.2 |
| 30 | 95 | 130.1 | 1.4 | 14.2 |
| 31 | 113 | 125.0 | 1.1 | 7.6 |
| **Overall** | **3,735** | **3,661.8** | **-** | **-** |

**Extended Data Table 3.** Localisation and LOD scores of markers in the QTL peak and confidence intervals for Cry1Ac resistance in the *Helicoverpa armigera* F2 intercross population.

| QTL | LG | Pos (cM) | LOD | P | Low Pos (cM) | High Pos (cM) |
| --- | --- | --- | --- | --- | --- | --- |
| Q1 | 5 | 62.76 | 6.48 | $3.29 \times 10^{-07}$ | 51.72 | 66.90 |
| Q2 | 12 | 2.76 | 8.80 | $1.57 \times 10^{-09}$ | 0.69 | 10.34 |

**Extended Data Table 4.**
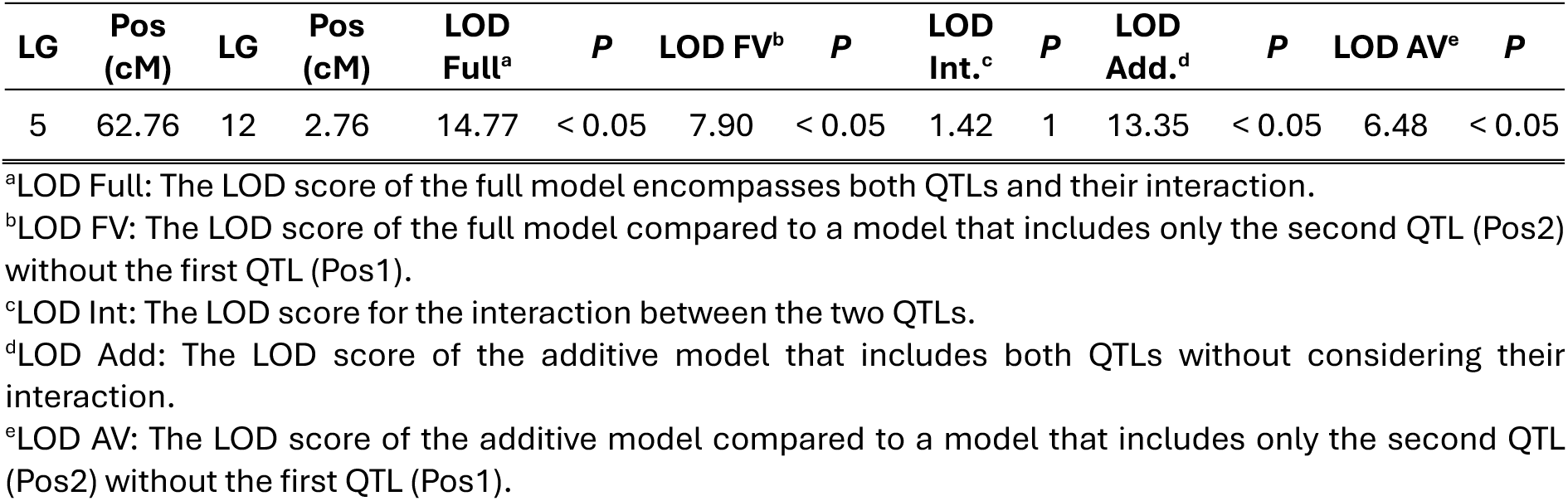
Summary table of the two-QTL genome scan to identify QTL interactions between the QTL involved in survival trait in *H*. *armigera* resistant to Cry1Ac.

**Extended Data Table 5.**
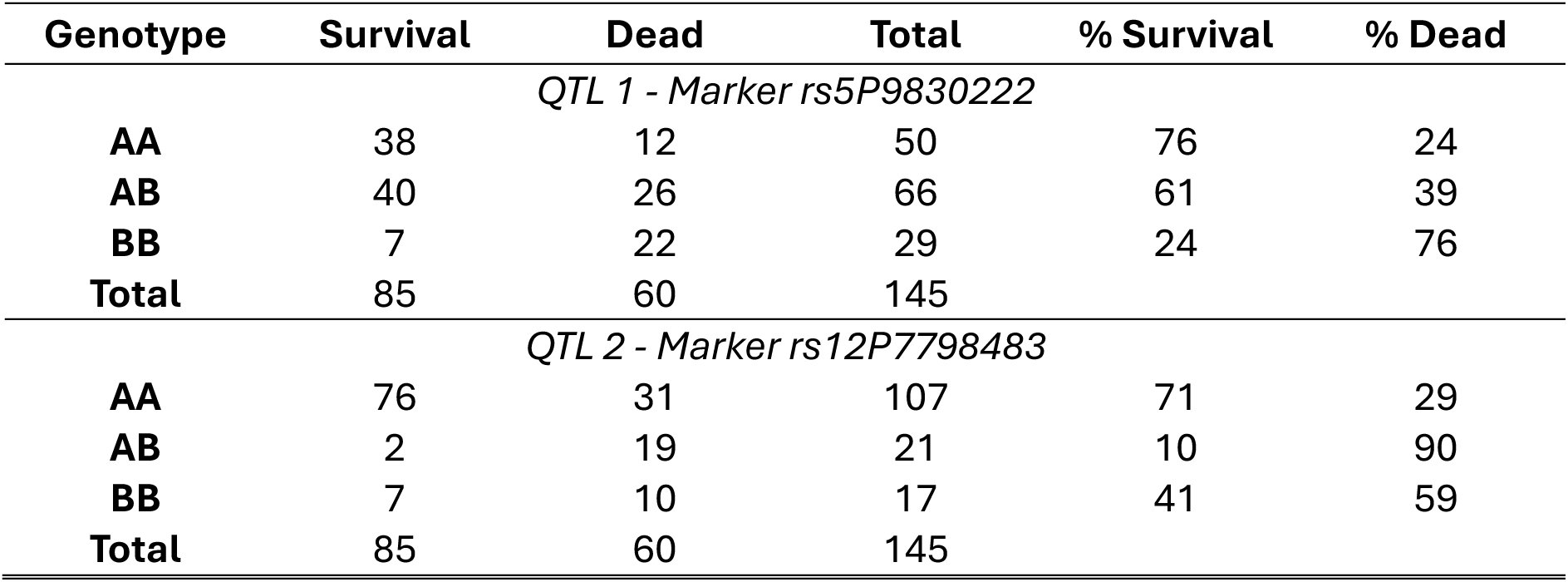
Genotypic frequencies of markers identified at QTL peaks associated with *Helicoverpa armigera* survival to the Cry1Ac protein.

| Genotype | Survival | Dead | Total | % Survival | % Dead |
| --- | --- | --- | --- | --- | --- |
| *QTL 1 - Marker rs5P9830222* |  |  |  |  |  |
| AA | 38 | 12 | 50 | 76 | 24 |
| AB | 40 | 26 | 66 | 61 | 39 |
| BB | 7 | 22 | 29 | 24 | 76 |
| Total | 85 | 60 | 145 |  |  |
| *QTL 2 - Marker rs12P7798483* |  |  |  |  |  |
| AA | 76 | 31 | 107 | 71 | 29 |
| AB | 2 | 19 | 21 | 10 | 90 |
| BB | 7 | 10 | 17 | 41 | 59 |
| Total | 85 | 60 | 145 |  |  |

**Extended Data Figure 1:**
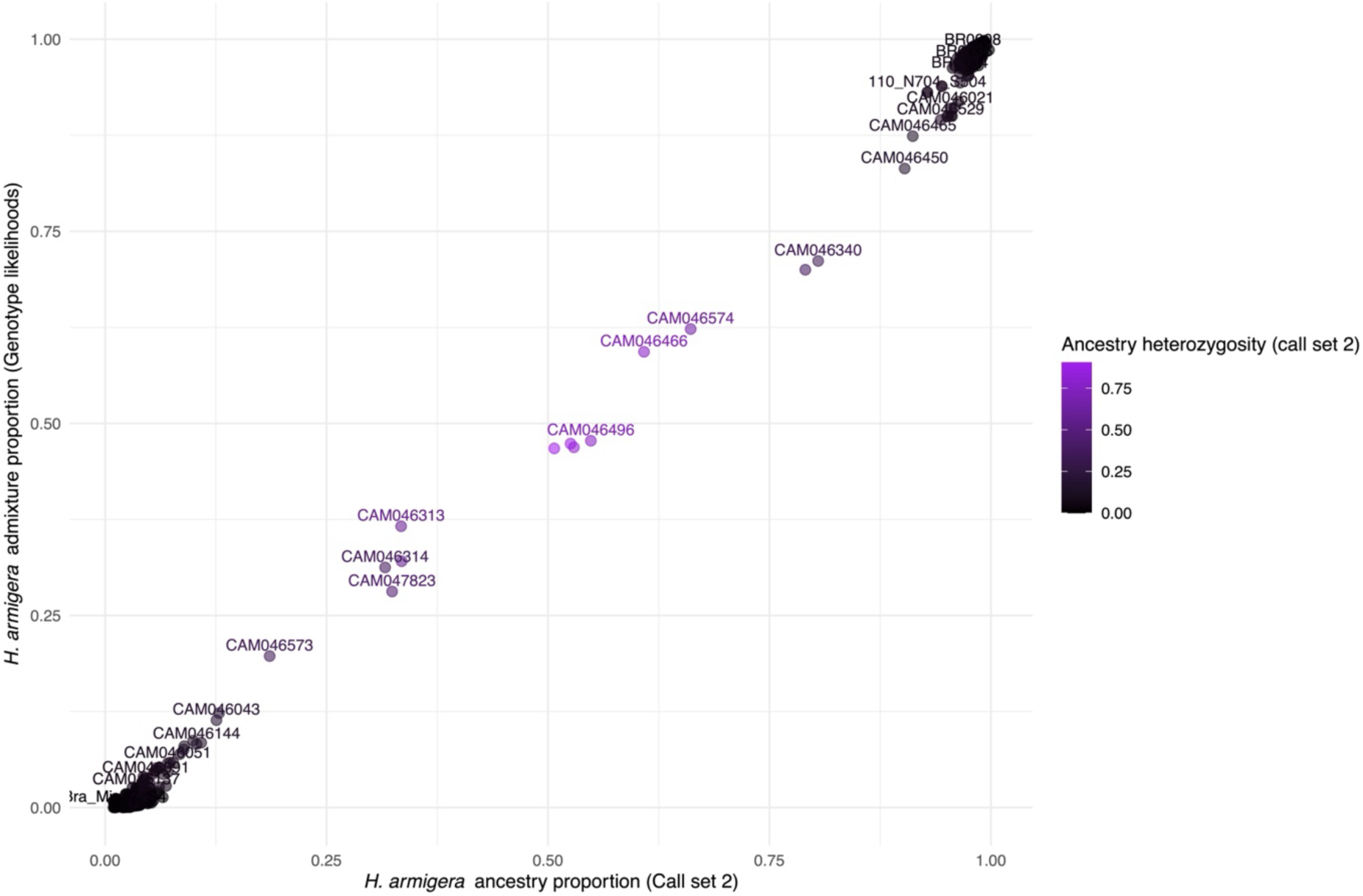
*H. armigera* ancestry proportion (AP) calculated using genotype calls and reference non-admixed *H. armigera* and *h. zea* (horizontal axis) against the ancestry proportion calculated using genotype likelihoods (vertical axis), with individual sample IDs labelled. Points and text are coloured by the ancestry heterozygosity calculated using genotype calls, which is maximised in early-generation hybrids (Figure 1).

**Extended Data Figure 2:**
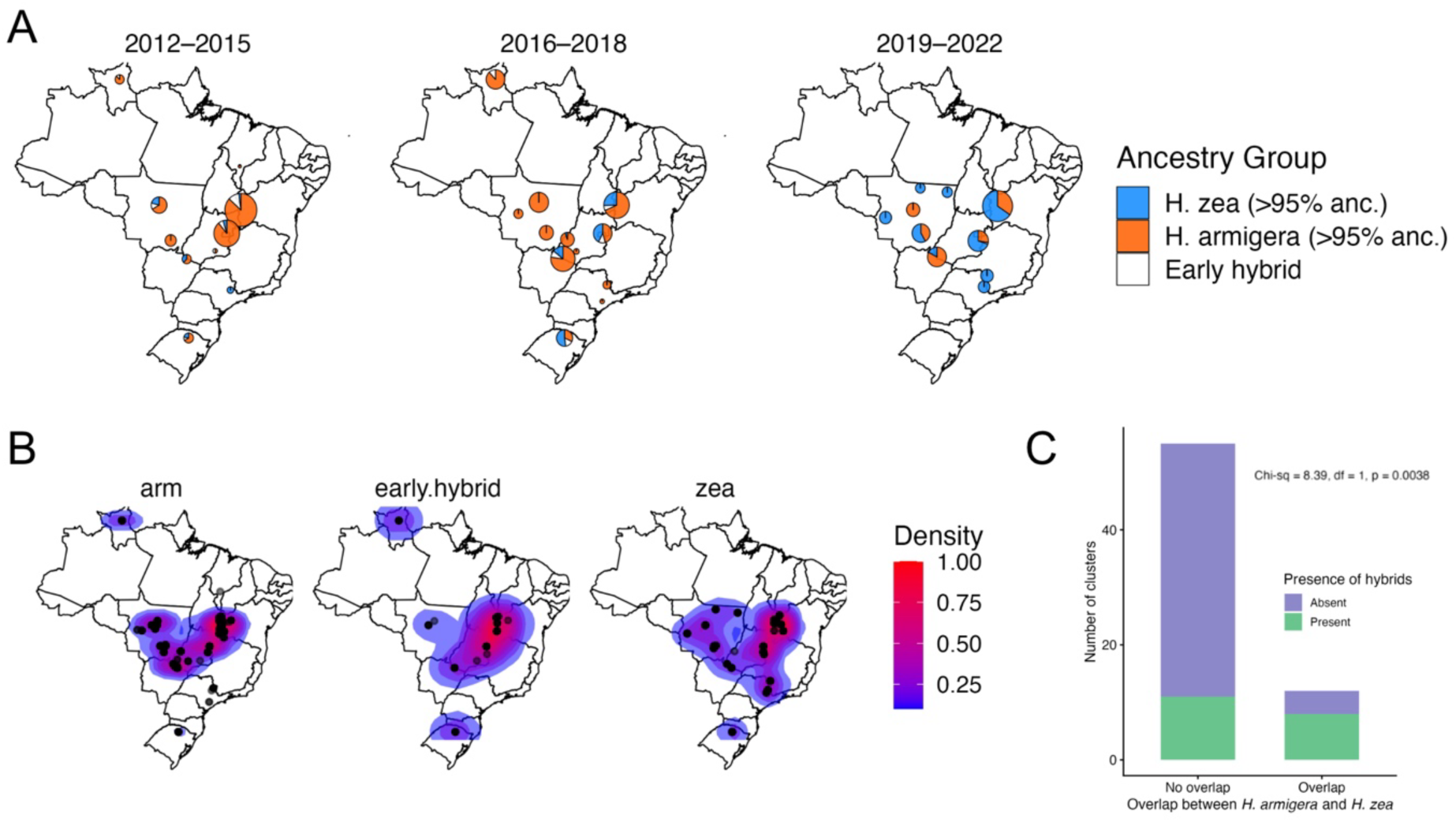
Spatial and temporal dynamics of Helicoverpa ancestry and early-generation hybrid occurrence in Brazil. **(A)** Proportion of individuals assigned to >95% H. armigera ancestry (orange), >95% H. zea ancestry (blue), or early generation hybrids (white) across Brazilian sampling sites grouped into three time periods: 2012–2015, 2016–2018, and 2019–2022 and in clusters whereby individuals are at a maximum of 100 km from each other. Pie chart size is proportional to sample size per locality. **(B)** Kernel density distributions showing spatial clustering of H. armigera, early hybrids, and H. zea across Brazil. Warmer colours indicate higher density of sampled individuals. **(C)** Proportion of sampling clusters where early hybrids are present or absent, split by regions with spatial overlap vs no spatial overlap of the two parental species. Hybrids are significantly more common in regions where the distributions of H. armigera and H. zea overlap (Chi–sq = 8.39, df = 1, p = 0.0038).

**Extended Data Figure 3:**
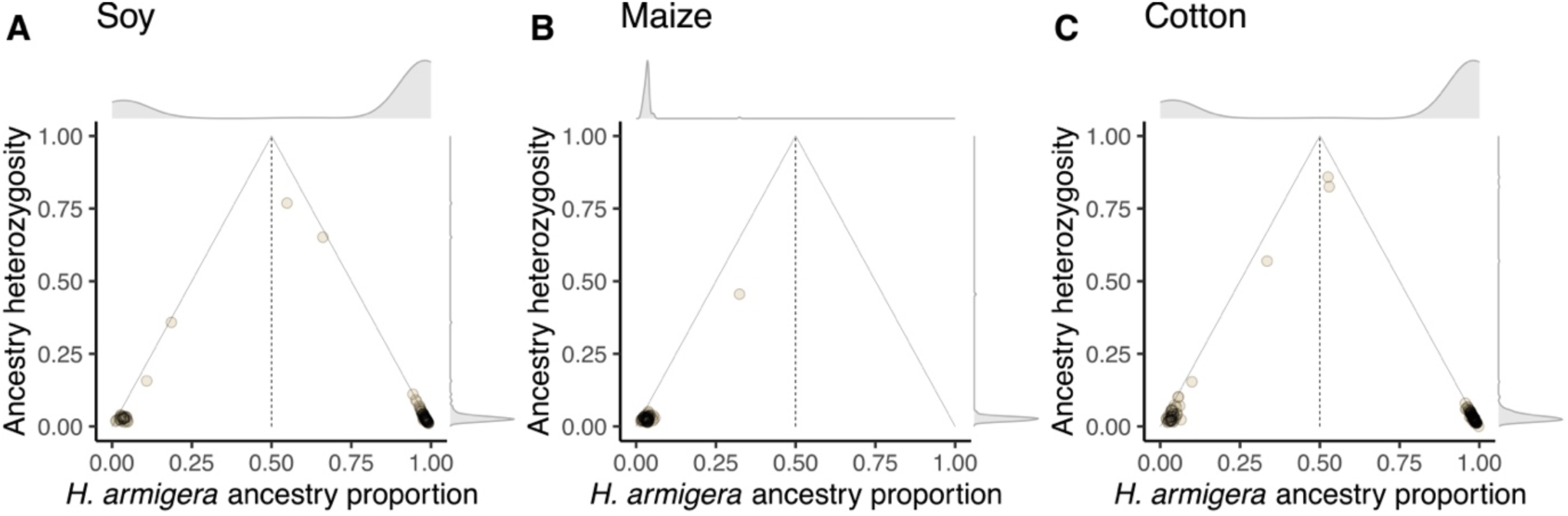
Individual ancestry proportion (AP) and ancestry heterozygosity (AH) across all timepoints and sample sites as for Figure 1C, for each crop from which individuals were sampled: maize, cotton, soybean (A-C respectively). Predominantly-*H. armigera* samples were collected on soybean and cotton, while predominantly-*H. zea* samples were collected on maize.

**Extended Data Figure 4:**
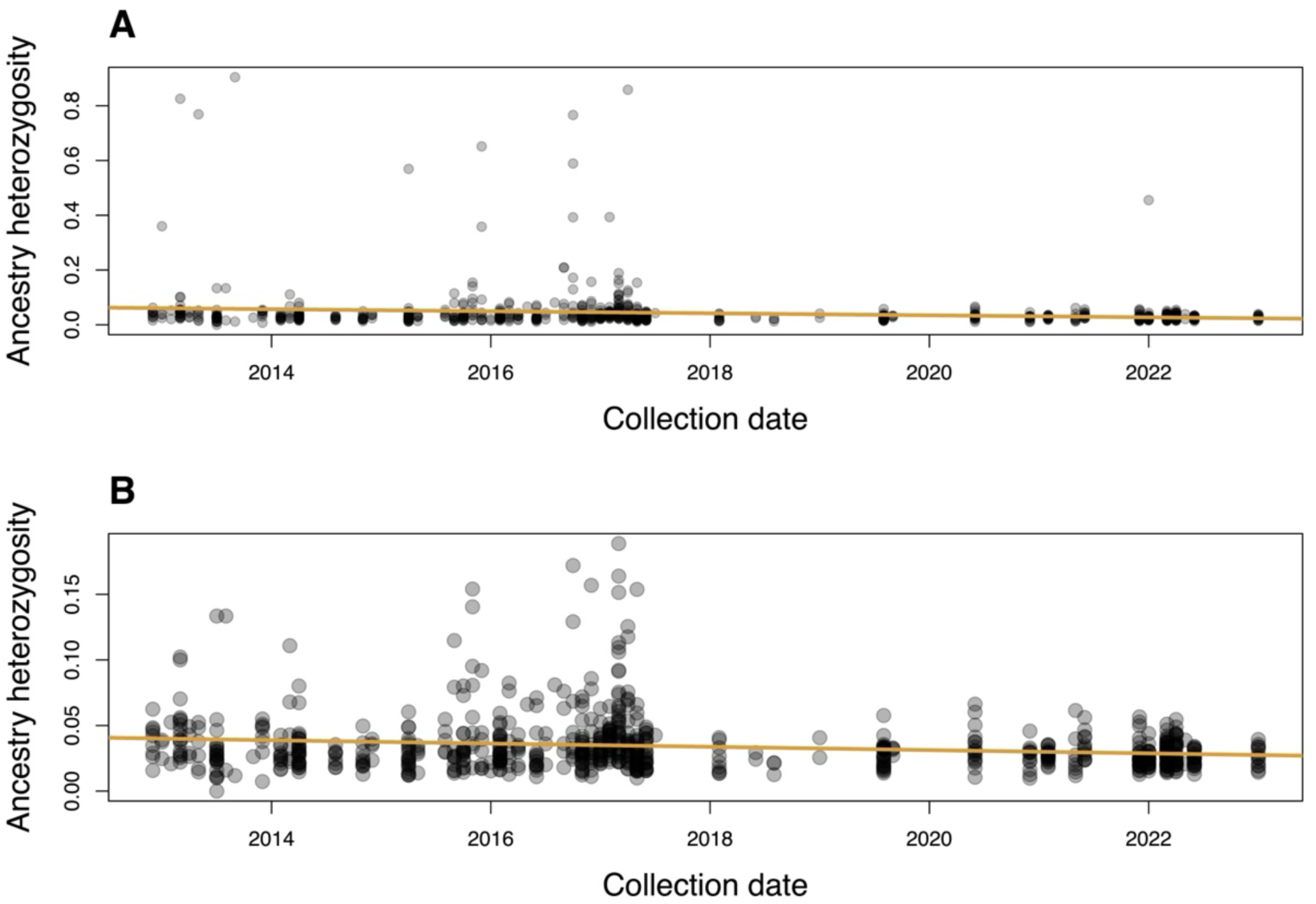
Individual ancestry heterozygosity vs collection date. Line shows a fitted linear model, which has a significantly negative gradient (Person’s product moment correlation: *p*<0.01, t = −4.9656, df = 746; correlation coefficient −0.177). **A**: all data shown. **B**: data shown for AH values <0.2.

**Extended Data Figure 5:**
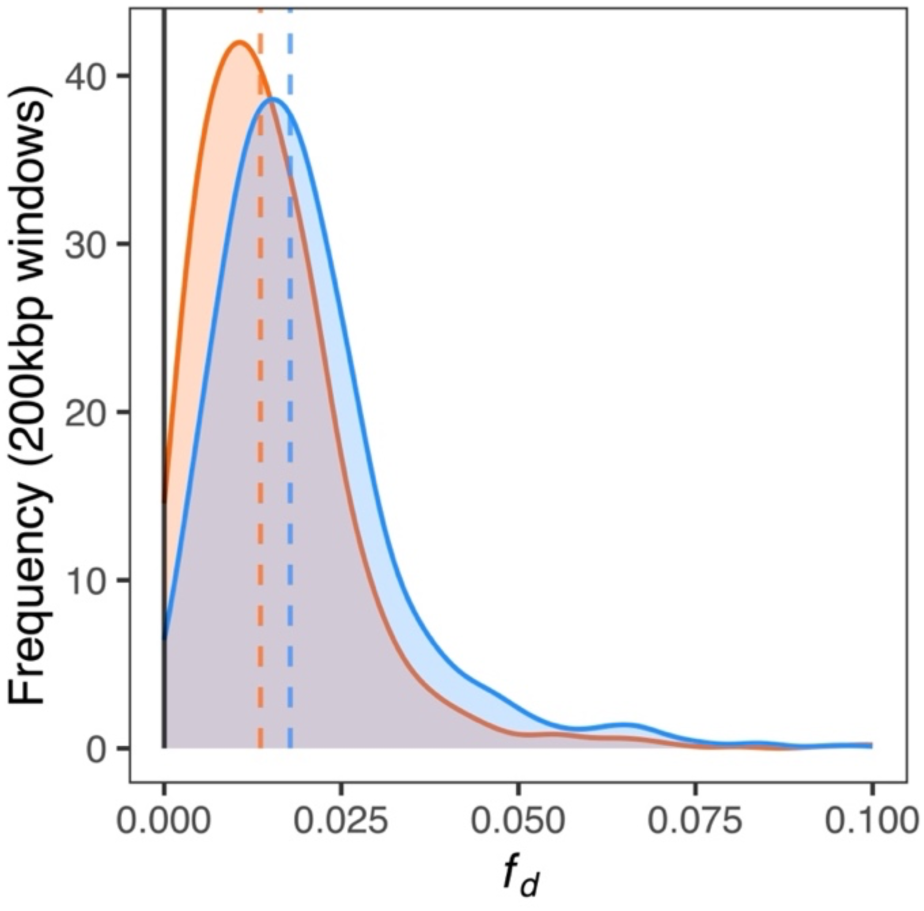
Distribution of *f_d_* calculated in 200kbp windows (horizontal axis limited to values between 0 and 0.1). Orange: gene flow from *H. armigera* to *H. zea* using the topology (((ZEA, ZEA_BRAZIL), ARM)), PUN); blue: gene flow from *H. zea* to *H. armigera* using the topology (((ARM, ARM_BRAZIL), ZEA)), PUN). Dashed lines indicate genome-wide medians: 0.0136 (*H. armigera* → *H. zea*), 0.0178 (*H. zea* → *H. armigera*).

**Extended Data Figure 6:**
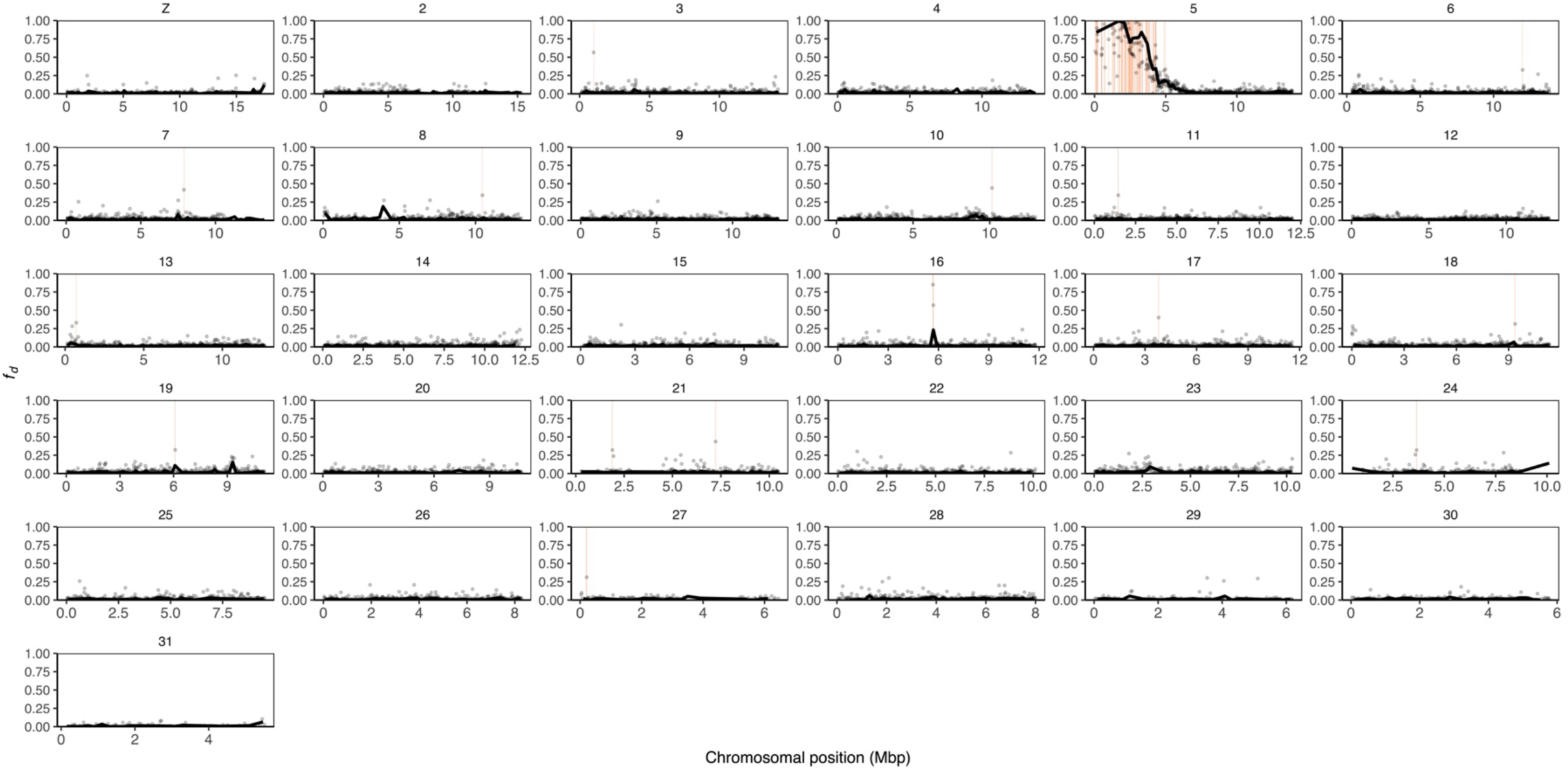
Genome-wide landscape of introgression from *H. armigera* to *H. zea* across all timepoints and locations. *f_d_* calculated in 200kbp windows (lines) and 20kbp windows (points) using the topology (((ZEA, ZEA_BRAZIL), ARM)), PUN). Highlighted windows indicate 20bkp windows in the upper first percentile of *f_d_* values (those equal to or greater than 0.310).

**Extended Data Figure 7:**
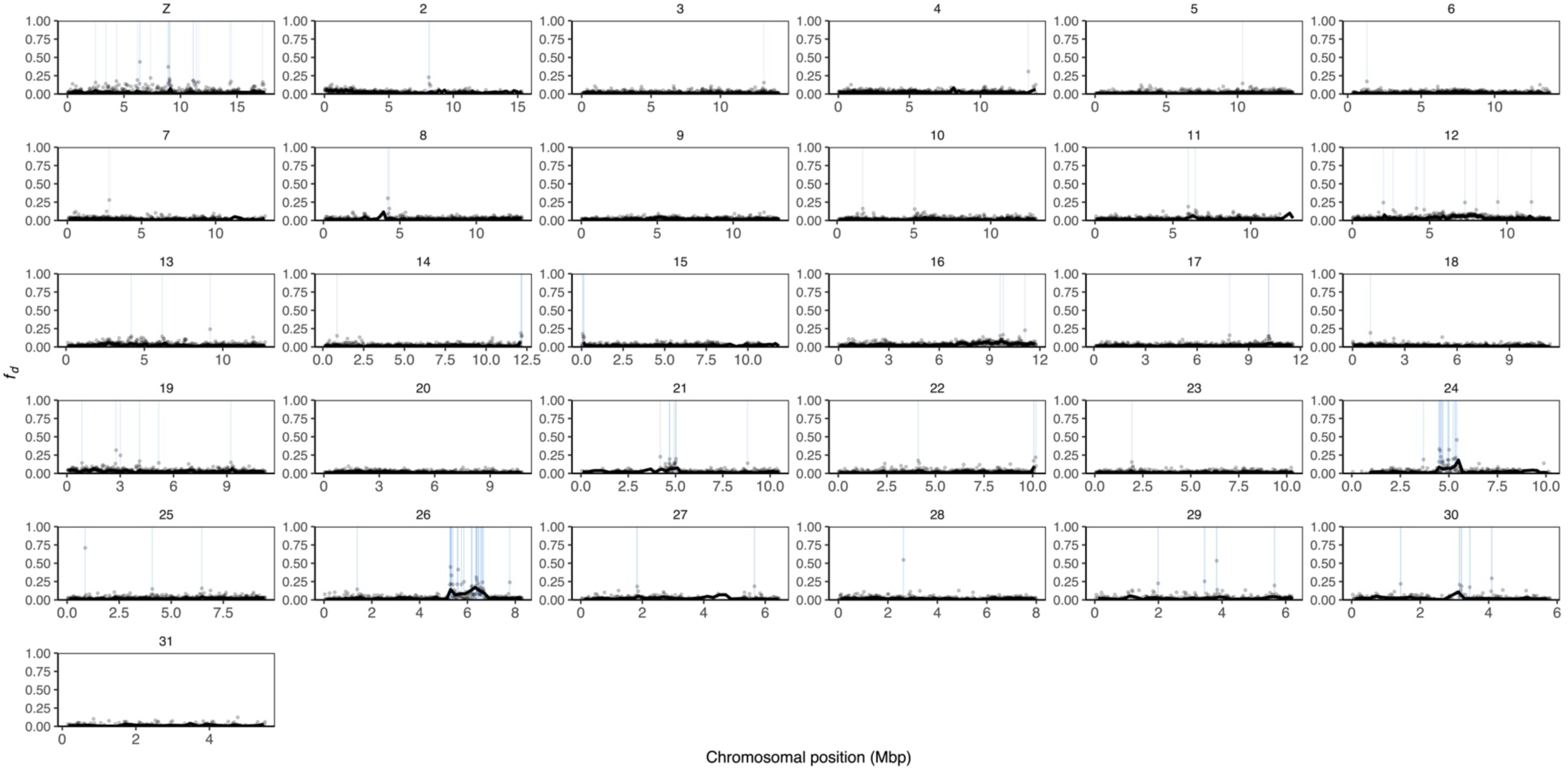
Genome-wide landscape of introgression from *H. zea* to *H. armigera* across all timepoints and locations. *f_d_* calculated in 200kbp windows (lines) and 20kbp windows (points) using the topology (((ZEA, ZEA_BRAZIL), ARM)), PUN). Highlighted windows indicate 20bkp windows in the upper first percentile of *f_d_* values (those equal to or greater than 0.139).

**Extended Data Figure 8:**
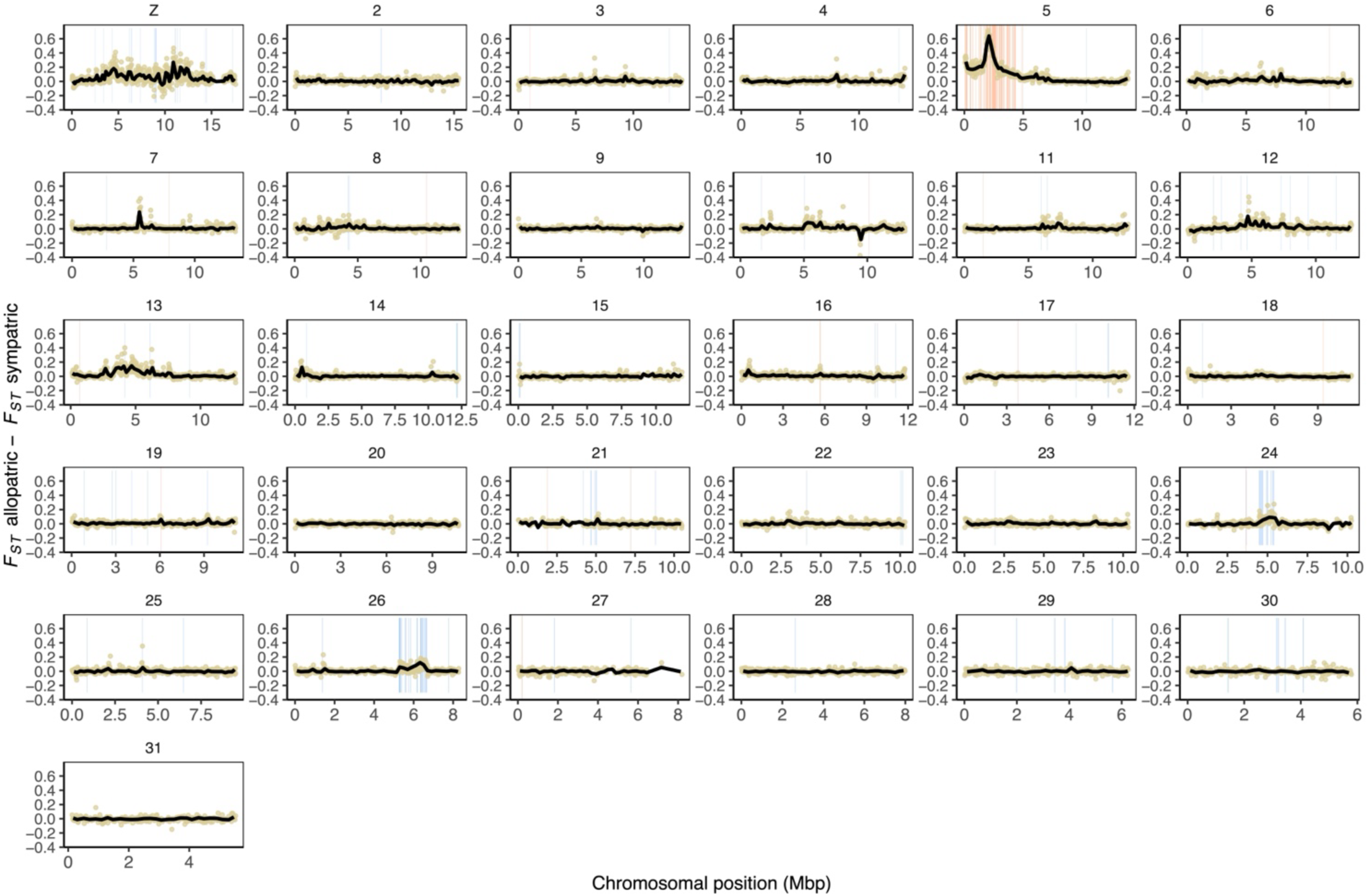
Local reductions in genetic differentiation in sympatry relative to allopatry, between *H. armigera* and *H. zea*. Genetic differentiation (*F_ST_*) was calculated in 200kbp windows with at least 500 informative sites (black lines) and 20kbp windows with at least 200 usable sites (points). “Allopatric” refers to the comparison of reference non-admixed *H. armigera* and *H. zea* in China and the United States respectively. “Sympatric” refers to the comparison of predominantly-*H. armigera* and predominantly-*H. zea* samples in Brazil, as defined by their ancestry proportion. Shaded regions indicate 20kbp windows with *f_d_* values in the upper first percentile (blue: *H. zea* → *H. armigera*; orange: *H. armigera* → *H. zea*).

**Extended Data Figure 9:**
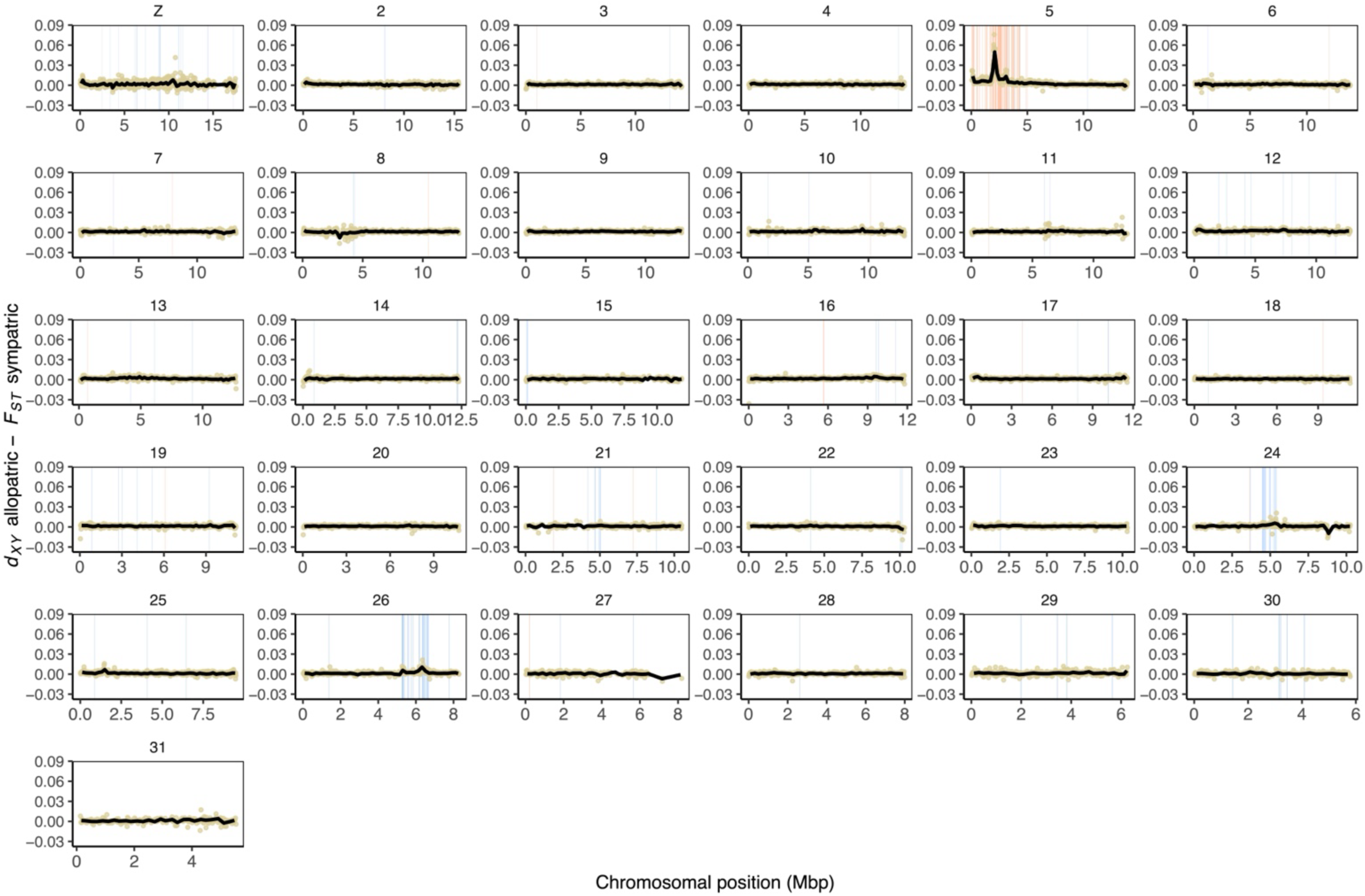
Local reductions in genetic divergence in sympatry relative to allopatry, between *H. armigera* and *H. zea*. Genetic divergence (*d_XY_*) was calculated in 200kbp windows with at least 500 informative sites (black lines) and 20kbp windows with at least 200 usable sites (points). “Allopatric” refers to the comparison of reference non-admixed *H. armigera* and *H. zea* in China and the United States respectively. “Sympatric” refers to the comparison of predominantly-*H. armigera* and predominantly-*H. zea* samples in Brazil, as defined by their ancestry proportion. Shaded regions indicate 20kbp windows with *f_d_* values in the upper first percentile (blue: *H. zea* → *H. armigera*; orange: *H. armigera* → *H. zea*).

**Extended Data Figure 10:**
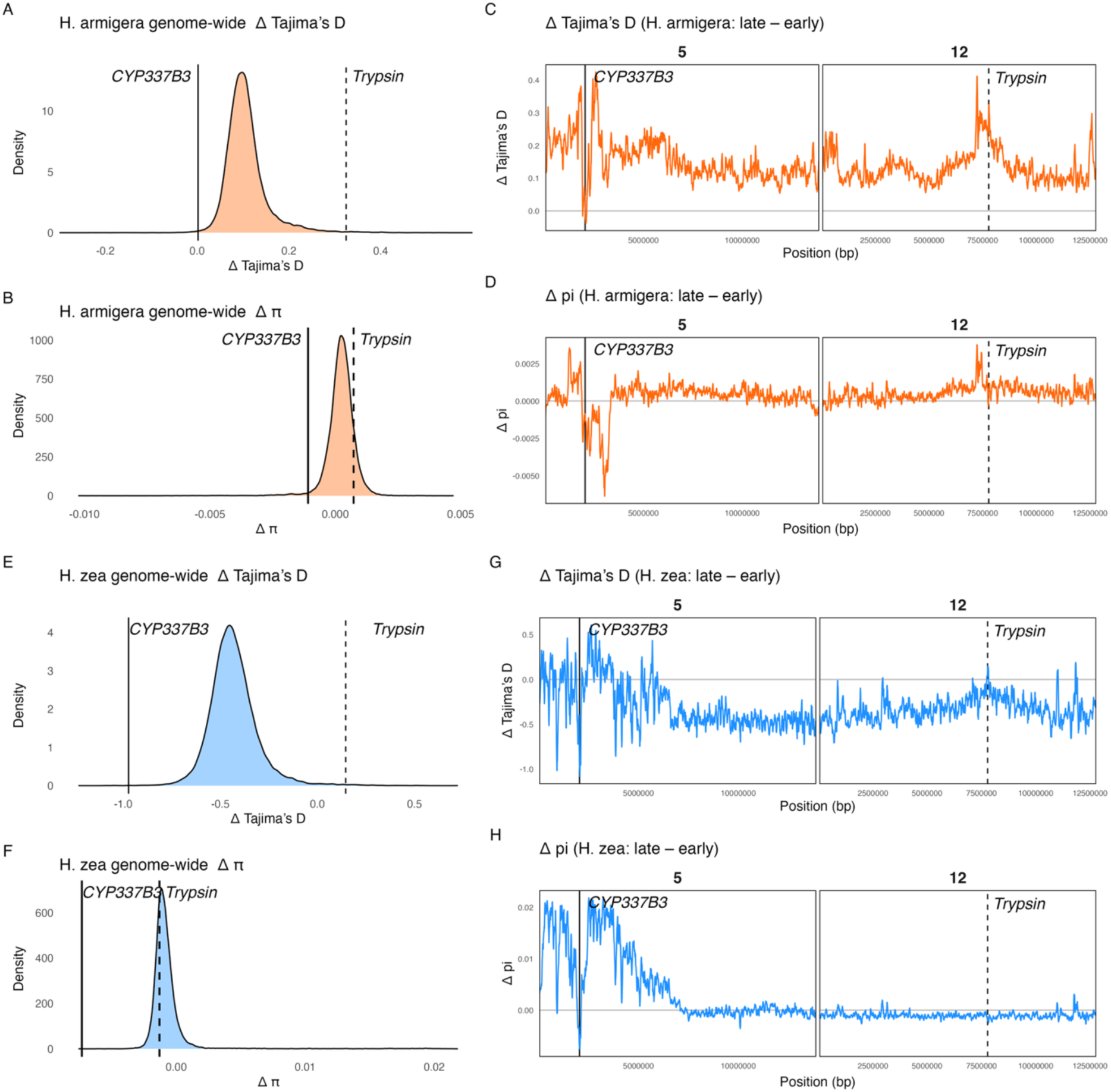
Genome-wide and locus-specific shifts in diversity statistics calculated with ANGSD between early and late collections of *Helicoverpa armigera* (A-D) and *H. zea* (E-H). (A, B) Genome-wide distributions of ΔTajima’s D (A, E) and Δπ (B, F) for *H. armigera* and *H. zea*, respectively (2012–2015 → 2019–2022). Vertical lines indicate the Δ values at CYP337B3 on chromosome 5 (solid) and the trypsin cluster on chromosome 12 (dashed). Chromosome-specific ΔTajima’s D (C, G) and Δπ (D, H) across chromosomes 5 and 12 in *H. armigera* and *H. zea*, with the positions of CYP337B3 and the trypsin cluster marked.

**Extended Data Figure 11:**
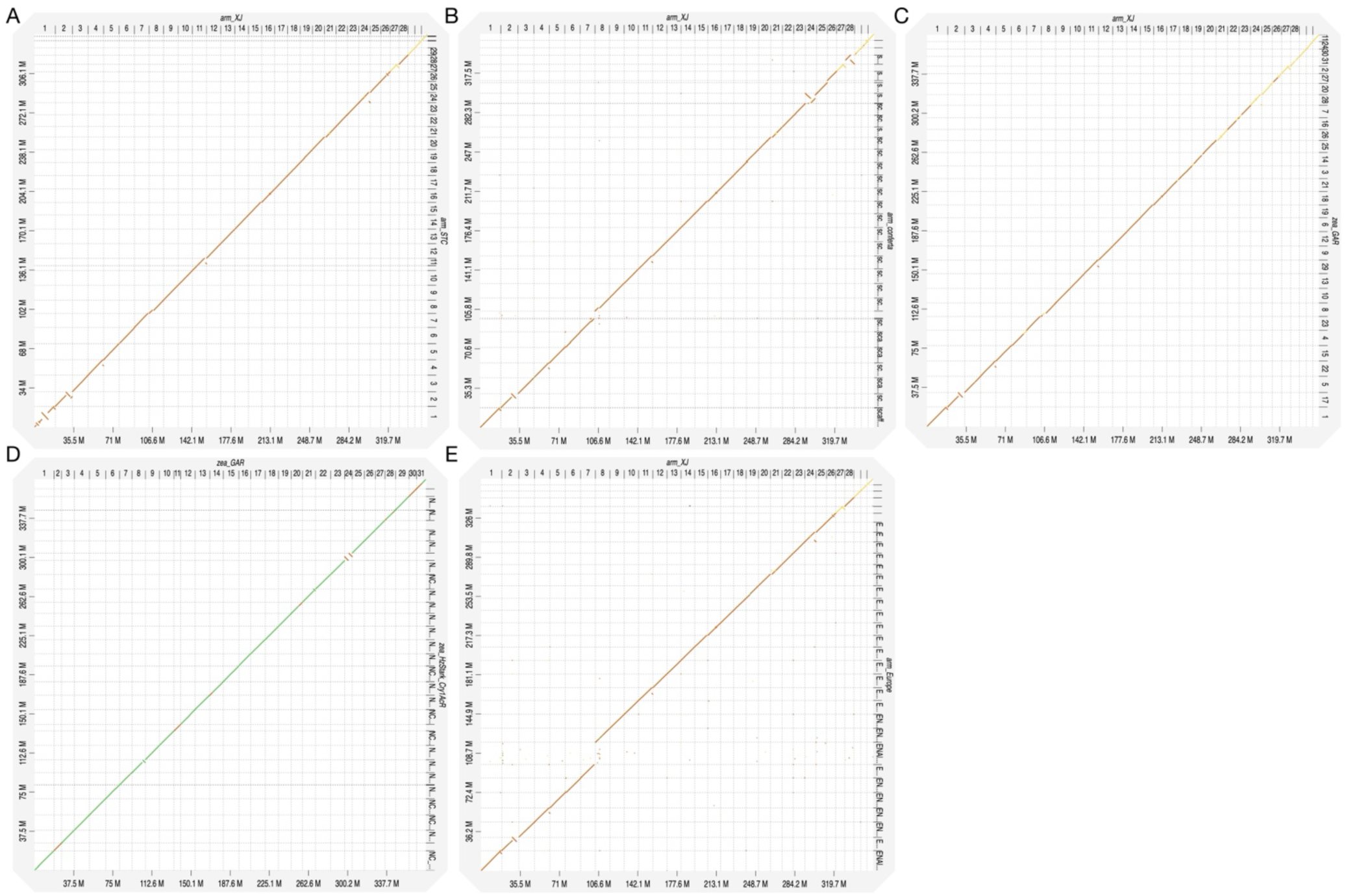
Synteny is generally highly conserved within and between *Helicoverpa armigera* and *Helicoverpa zea*. Axes show scaffold positions of all 31 homologous chromosomes, where “1” corresponds to the Z chromosome in the arm_XJ reference genome used here. Grid lines delineate scaffolds. Colour corresponds to genetic divergence (green=low, yellow=medium, orange=high). **(A)** Comparison of the reference genome used here, derived from a sample collected in north-western China (horizontal axis) and a reference genome of a sample collected from a distinct *H. armigera* population in southern China (vertical axis) ^34^ **(B)** Comparison of the *H. armigera* reference genome used here (horizontal axis) and a reference genome generated for *H. armigera conferta*, collected in Australia (vertical axis) ^29^. **(C)** Comparison of the *H. armigera* reference genome used here (horizontal axis) and the *H. zea* reference genome generated using the GA-R strain (vertical axis) ^27^. **(D)** Comparison of the *H. zea* GA-R reference genome (vertical axis) and the *H. zea* HzStark_Cry1AcR strain ^28^. **(E)** Comparison of the *H. armigera* reference genome used here (horizontal axis) and a *H. armigera* reference genome generated using a sample collected in Europe (vertical axis) ^30^.

**Extended Data Figure 12:**
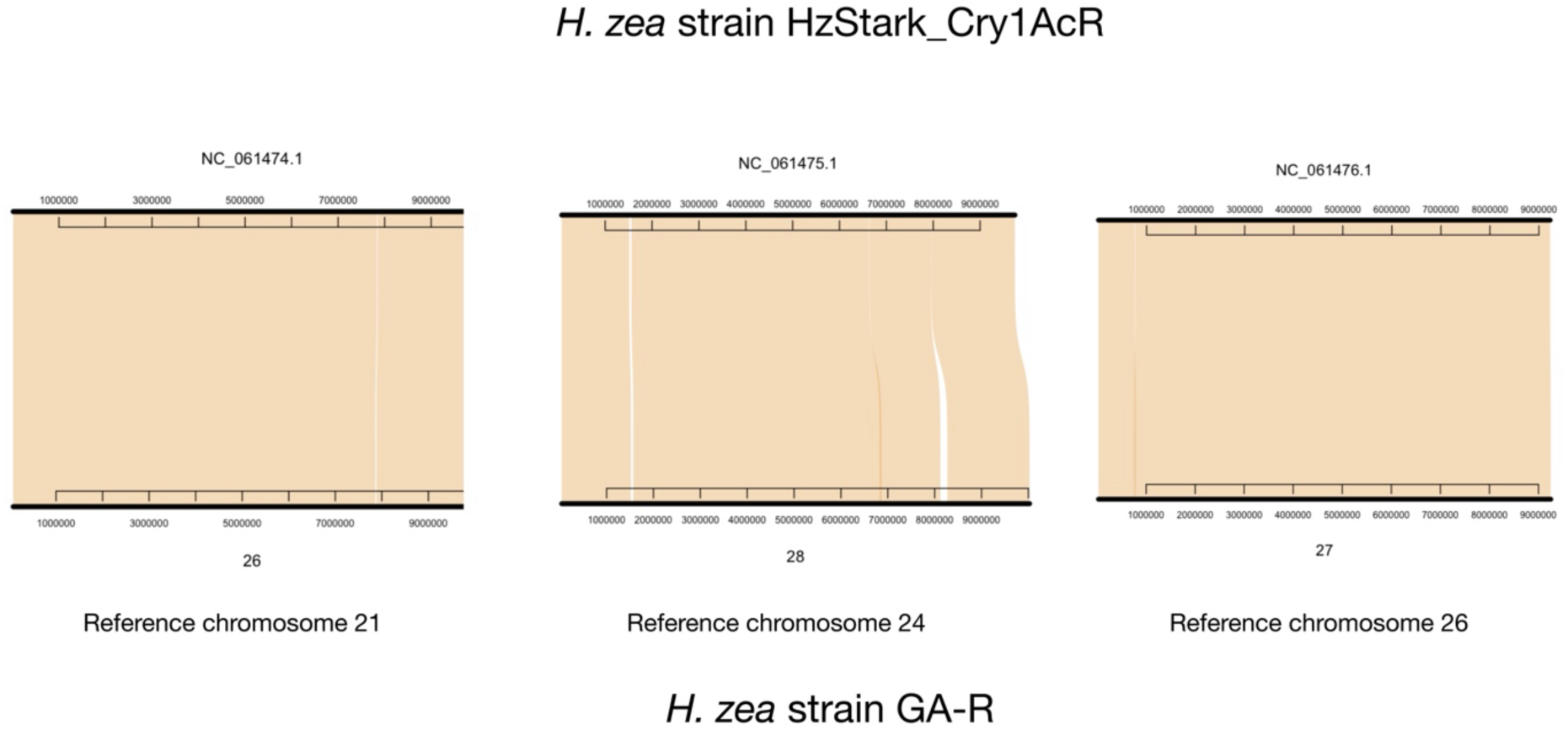
No inversion polymorphism between *H. zea* reference assemblies at the three chromosomes of interest. GA-R chromosomes 26, 28 and 27 correspond to chromosomes 21, 24 and 26 in the reference genome used here. Shaded regions show nucleotide alignments between the GA-R reference genome (lower scaffolds) ^27^ and the *H. zea* HzStark_Cry1AcR strain (upper scaffolds) ^28^.

**Extended Data Figure 13:**
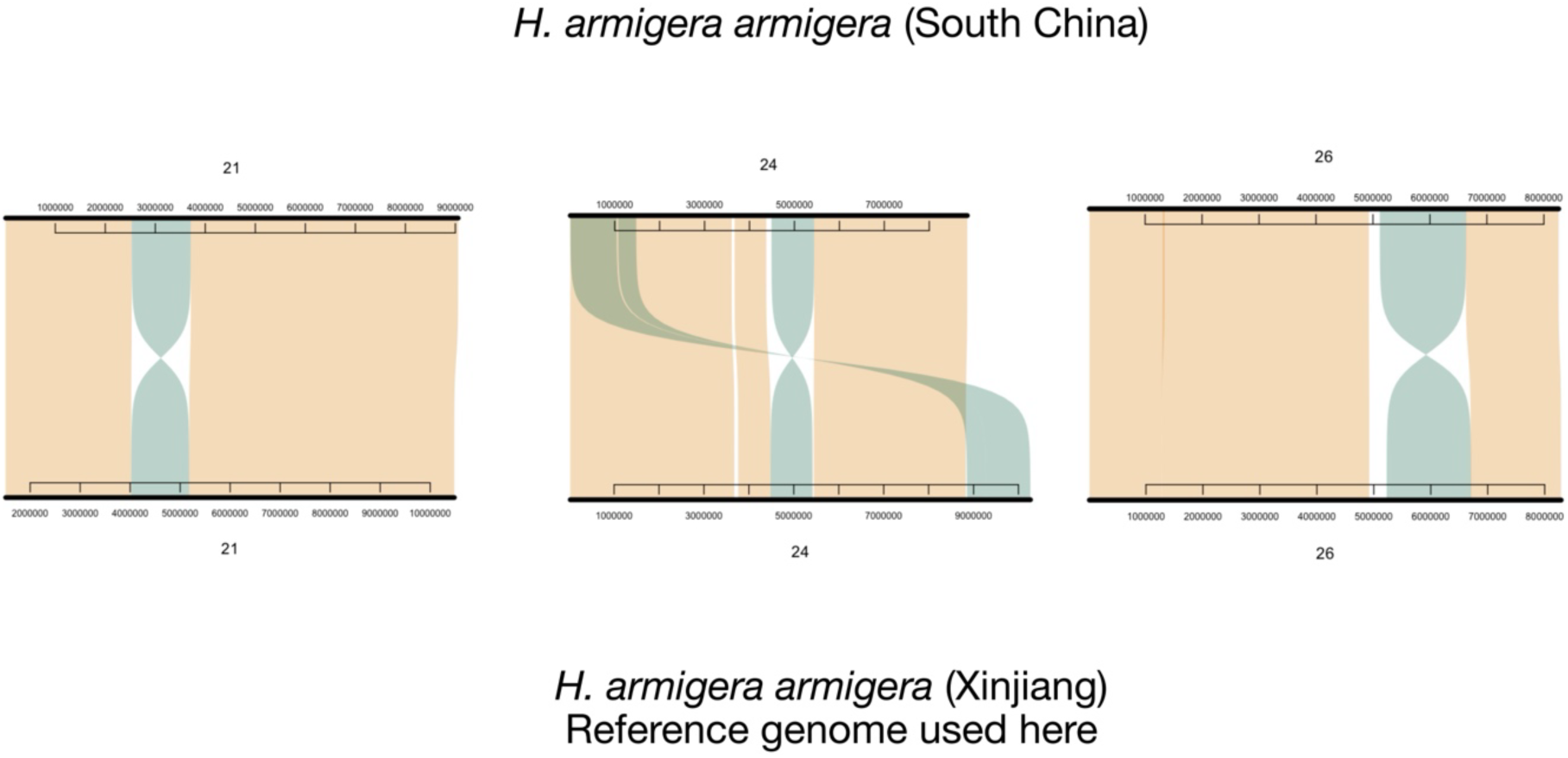
Inversions polymorphic between *H. armigera* and *H. zea* are also polymorphic between distinct *H. armigera* lineages. Shaded regions show nucleotide alignments between the reference genome used here (lower scaffolds) and a reference assembly generated from a sample collected in South China (upper scaffolds) ^34^.

**Extended Data Figure 14:**
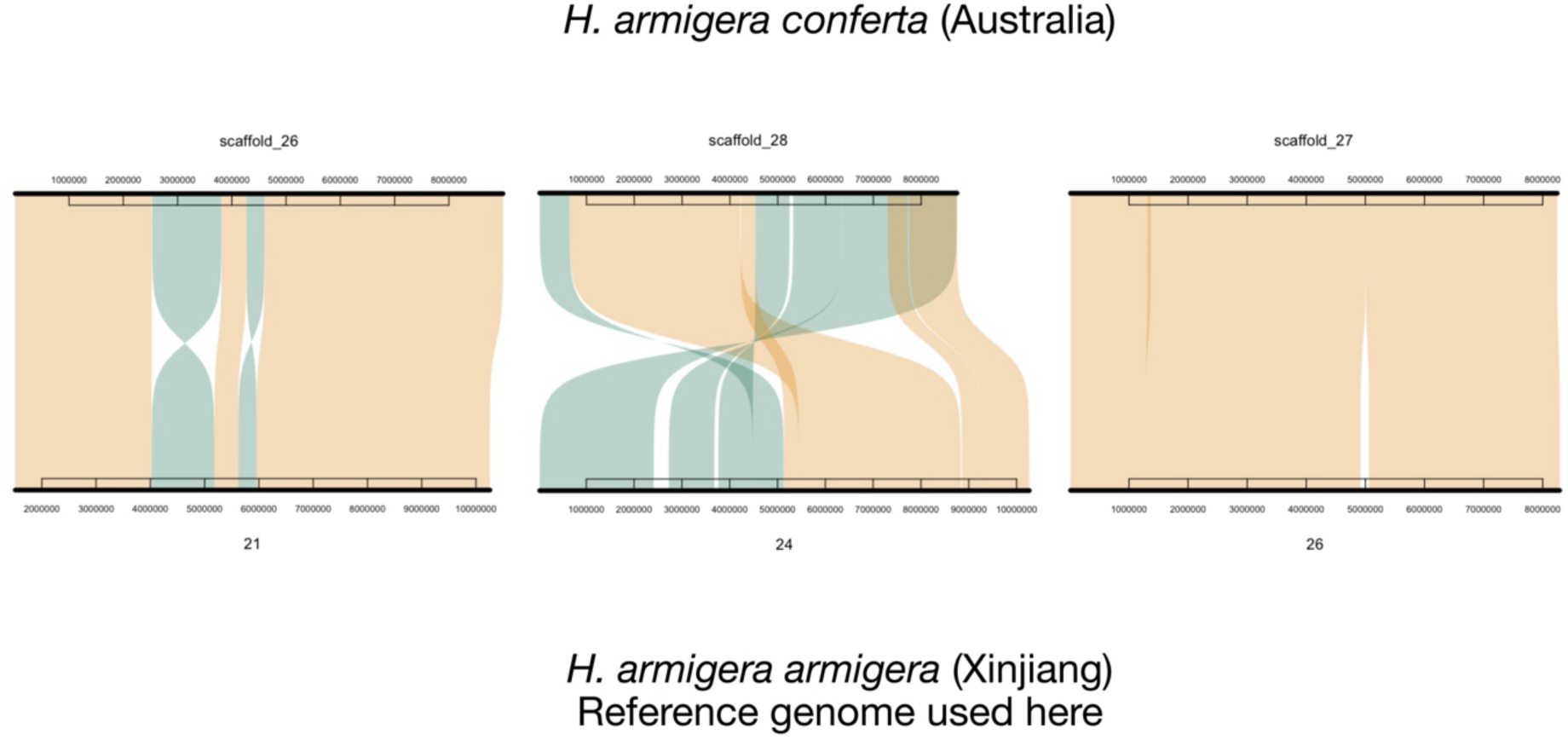
On chromosome 26, inversions polymorphic between *H. armigera* and *H. zea* are not polymorphic between *H. armigera armigera* and *H. armigera conferta*. The other two inversions of interest are polymorphic within *H. armigera*. Shaded regions show nucleotide alignments between the reference genome used here (lower scaffolds) and a *H. armigera conferta* reference assembly ^29^.

**Extended Data Figure 15:**
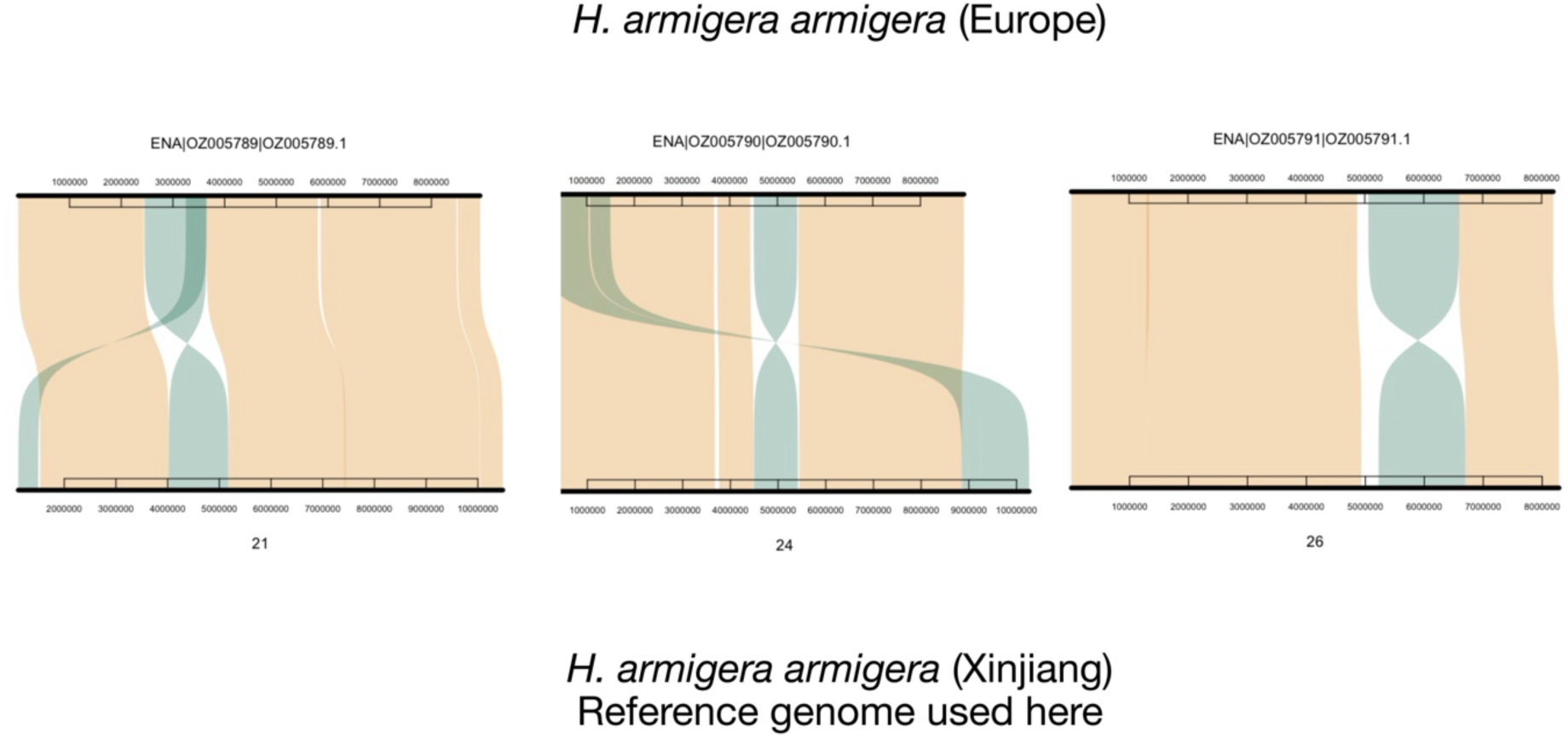
Inversions polymorphic between *H. armigera* and *H. zea* are also polymorphic between *H. armigera* populations. Shaded regions show nucleotide alignments between the reference genome used here (lower scaffolds) and a reference assembly generated from a sample collected in Europe (upper scaffolds) ^30^

**Extended Data Figure 16:**
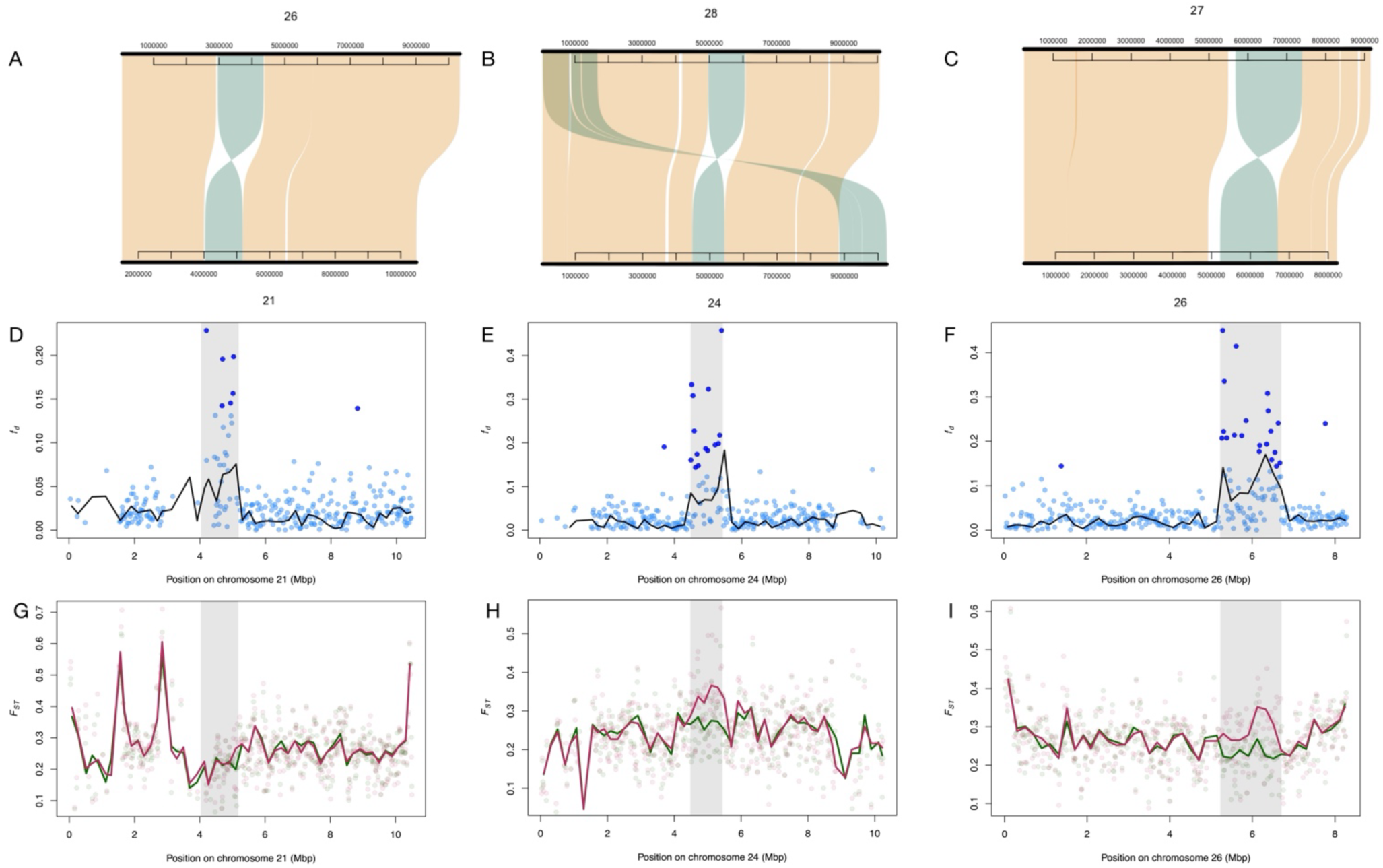
Introgression of three *H. zea* inversions into *H. armigera*. All three inversions arose within *H. armigera*, so the introgressed *H. zea* haplotype is the ancestral orientation. **(A-C)** Synteny maps comparing the *H. armigera* reference genome used here (bottom scaffold) to the *H. zea* GA-R reference genome ^27^ for chromosomes 21, 24 and 26 respectively. Scaffold numbers indicate homologous chromosomes in the two assemblies (upper: *H. zea*, lower: *H. armigera*). **(D-F)** *f_d_* (*H. zea* → *H. armigera*) calculated in non-overlapping 200kbp windows (lines) and 20kbp non-overlapping windows (points). Dark coloured points indicate those in the upper first percentile of *f_d_* values genome-wide. Shaded regions indicate the bounds of the inversion identified in A. **(G-I)** *F_ST_* calculated between allopatric non-admixed *H. armigera* and *H. zea* (maroon) and sympatric Brazilian *H. armigera* and *H. zea* (green). Lines: 200kbp windows, points: 20kbp windows.

**Extended Data Figure 17:**
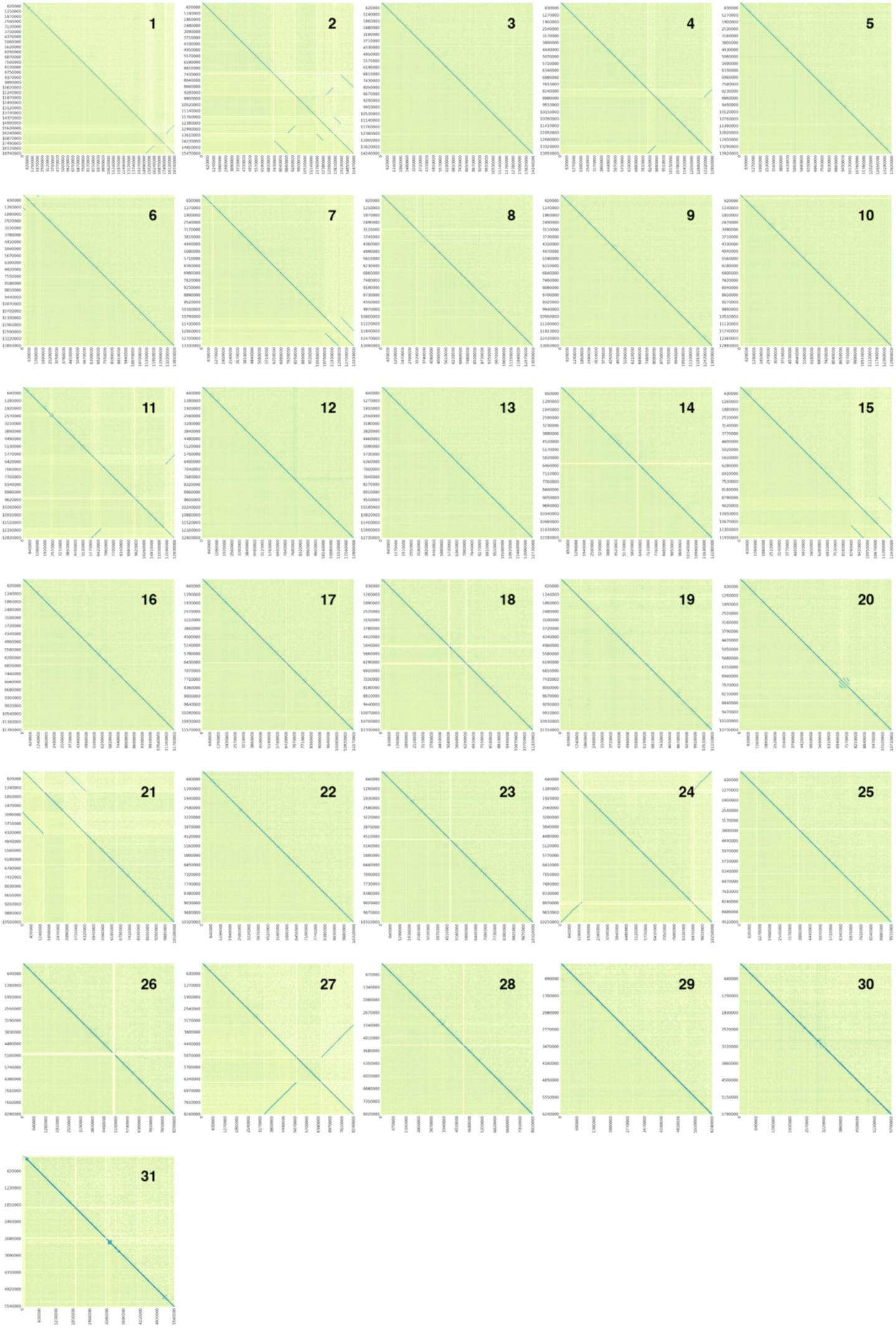
Genome-wide barcode-sharing heatmaps used to detect structural variants in *H. zea* and *H. armigera*. Heatmaps show Wrath-derived barcode-sharing matrices across all chromosomes, calculated from linked-read BAM files using 10-kb non-overlapping windows. For each chromosome, the upper triangular matrix represents the *H. zea* dataset and the lower triangular matrix represents the *H. armigera* dataset. Colours reflect the Jaccard index of shared barcodes between window pairs, with darker regions indicating stronger barcode similarity. This method captures local deviations from the expected decay of barcode sharing with distance and highlights putative structural variants. Chromosome numbers are displayed in the upper right of each panel. Heatmaps correspond to the same matrices used by Wrath’s SV-calling module (–l), which applies z-score outlier detection and a double exponential decay model to identify structural rearrangements.

**Extended Data Figure 18:**
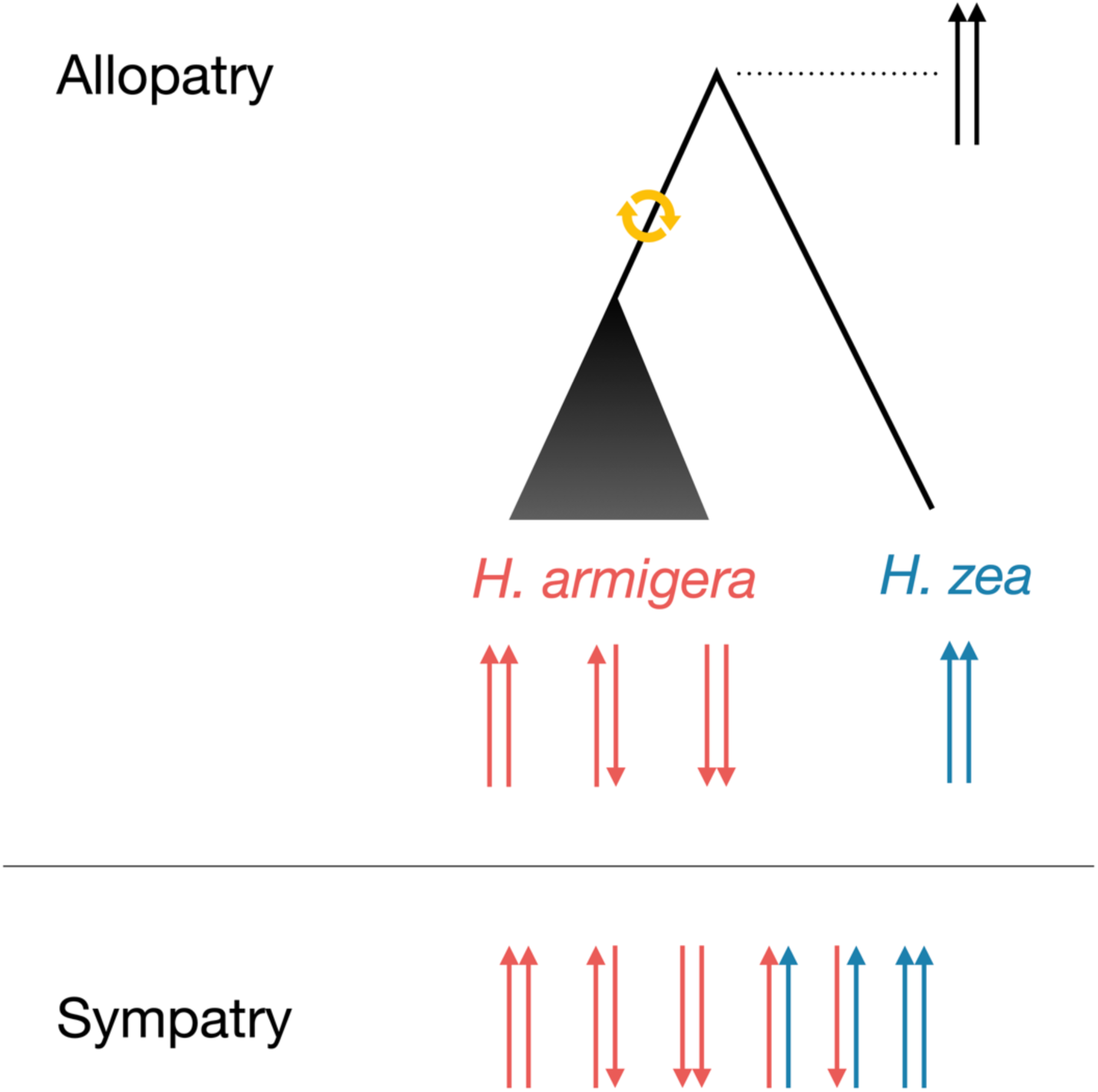
Proposed scenario for all three inversion polymorphisms on chromosomes 21, 24 and 26, based on the most parsimonious explanation for inversion polymorphisms. An inversion mutation occurred within *H. armigera*, resulting in a derived inverted allele that remains polymorphic across divergent *H. armigera* lineages. The mutation occurred in a population ancestral to multiple present-day *H. armigera* lineages, or arose in one *H. armigera* lineage and spread to others. There are therefore three possible genotypes in allopatric *H. armigera* and one possible genotype in allopatric *H. zea*. Upon secondary sympatry in Brazil, there are six possible genotypes. Based on the available data I cannot exclude the possibility that the inversion was present in the ancestor to *H. armigera* and *H. zea*, but was lost in *H. zea*.

**Extended Data Figure 19:**
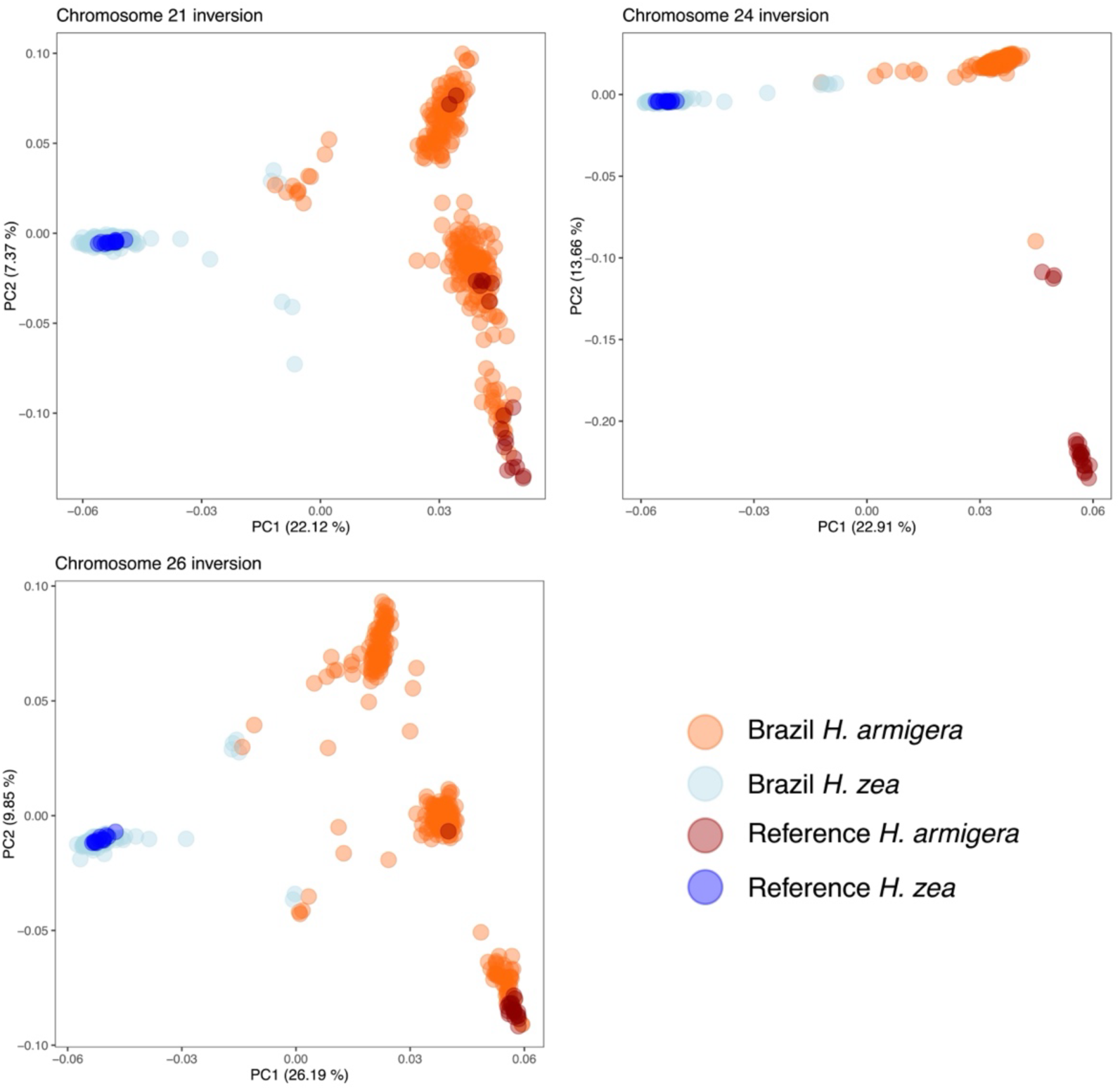
PCA in in inversion regions confirms introgression from *H. zea* to *H. armigera*. Point colour corresponds to ancestry, as defined by the likelihood-based ancestry proportion. Unlike the pattern outside of inverted regions in which individuals cluster by ancestry proportion, in inversion regions individuals fall into three major clusters with up three rarer intermediate clusters. This is consistent with expectations under the proposed evolutionary scenario for all three inversions (Extended Data Fig. 18). Heterozygous individuals are overwhelmingly *H. armigera* samples, consistent with the introgression signal.

**Extended Data Figure 20:**
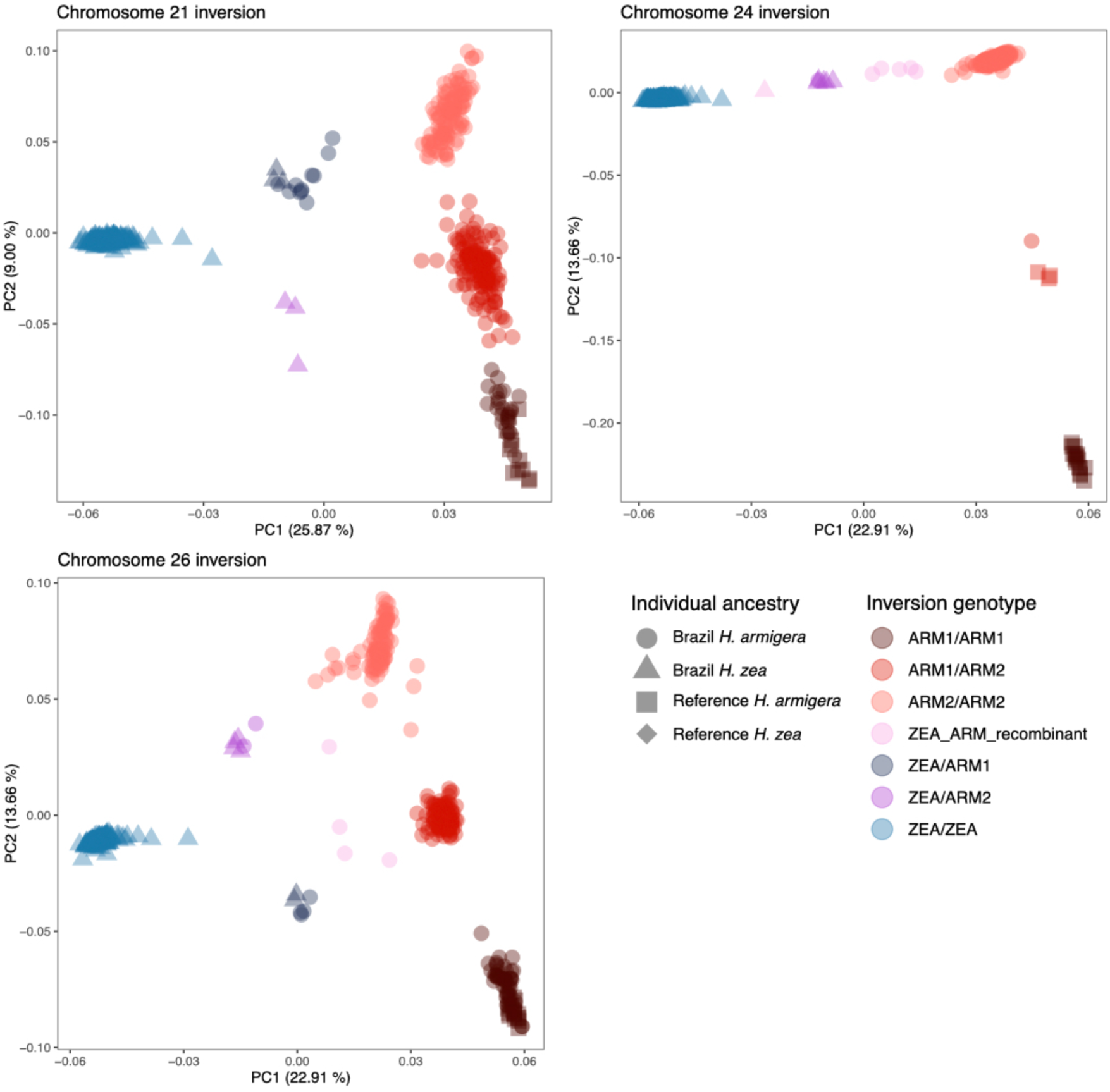
Putative inversion genotypes assigned using PCA clusters. Shape corresponds to ancestry, colour corresponds to genotype assignment. This approach distinguishes between two possible *H. armigera* alleles, but does not assign which is ancestral or derived. This is because of the significant degree of genetic differentiation between the three putative homozygous genotypes. Unlike the pattern outside of inverted regions in which individuals cluster by ancestry proportion, in inversion regions individuals fall into three major clusters with up three rarer intermediate clusters. This fits expectations under the proposed evolutionary scenario for all three inversions. Individual genotype assignments are reported in Extended Data Table 1. Recombinant genotype assignments fall between genotype categories, representing possible cases of recombination or gene conversion.

**Extended Data Figure 21.**
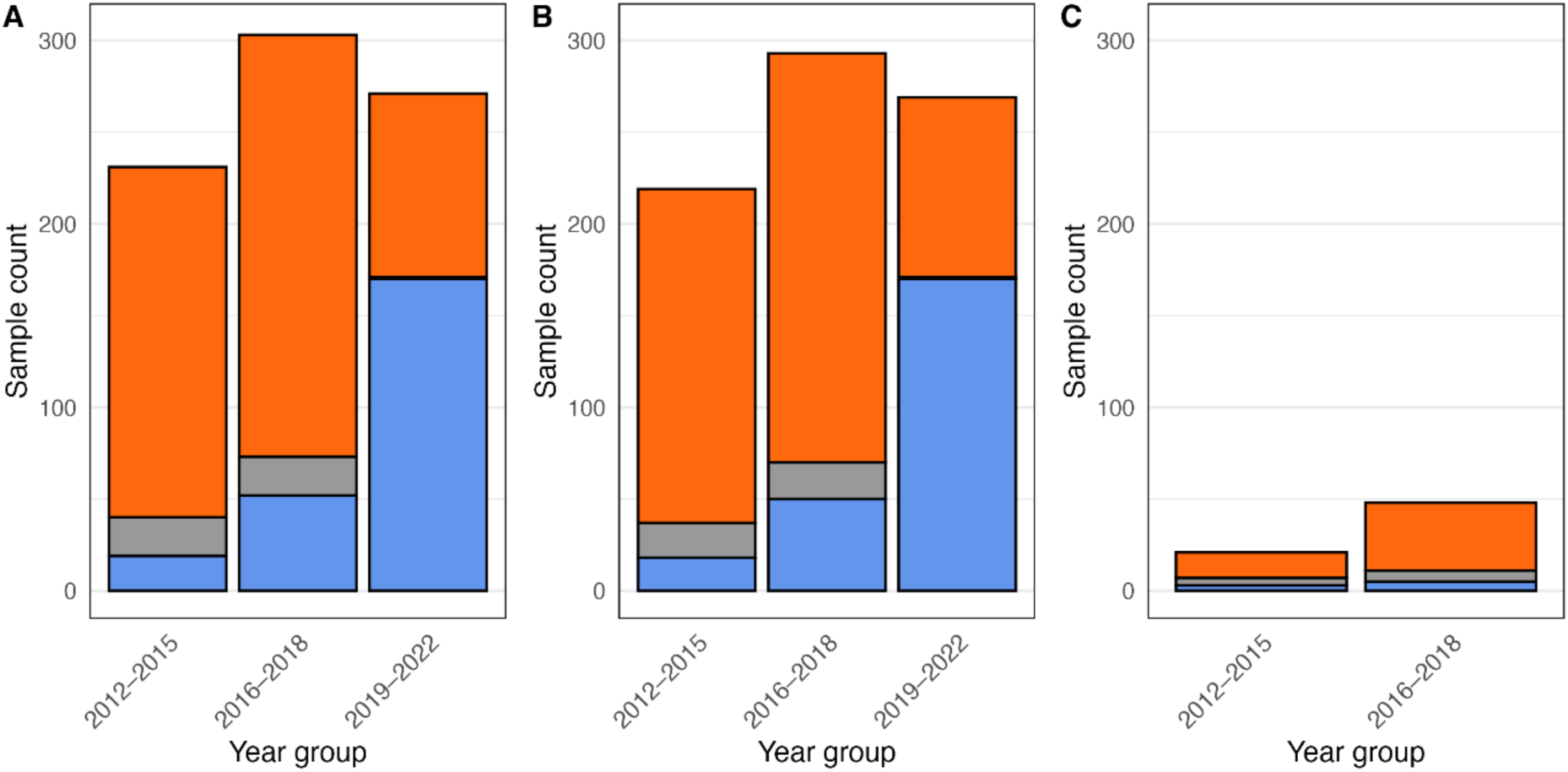
Population ancestry composition among Brazilian Helicoverpa samples across three genomic datasets used for these analyses. (A) Genotype-likelihood dataset (n = 820 individuals), (B) variant-calling dataset (n = 796), and (C) haplotagging dataset (n = 93). Bars show absolute counts of individuals per year-group, coloured by ancestry group as determined by PCAngsd (armigera-like, zea-like, and early generation hybrid categories). All datasets include only Brazilian and newly sequenced individuals; additional published samples were excluded for consistency across panels. All individuals in the variant-calling dataset (B) are included in genotype-likelihood dataset (A).

**Extended Data Figure 22:**
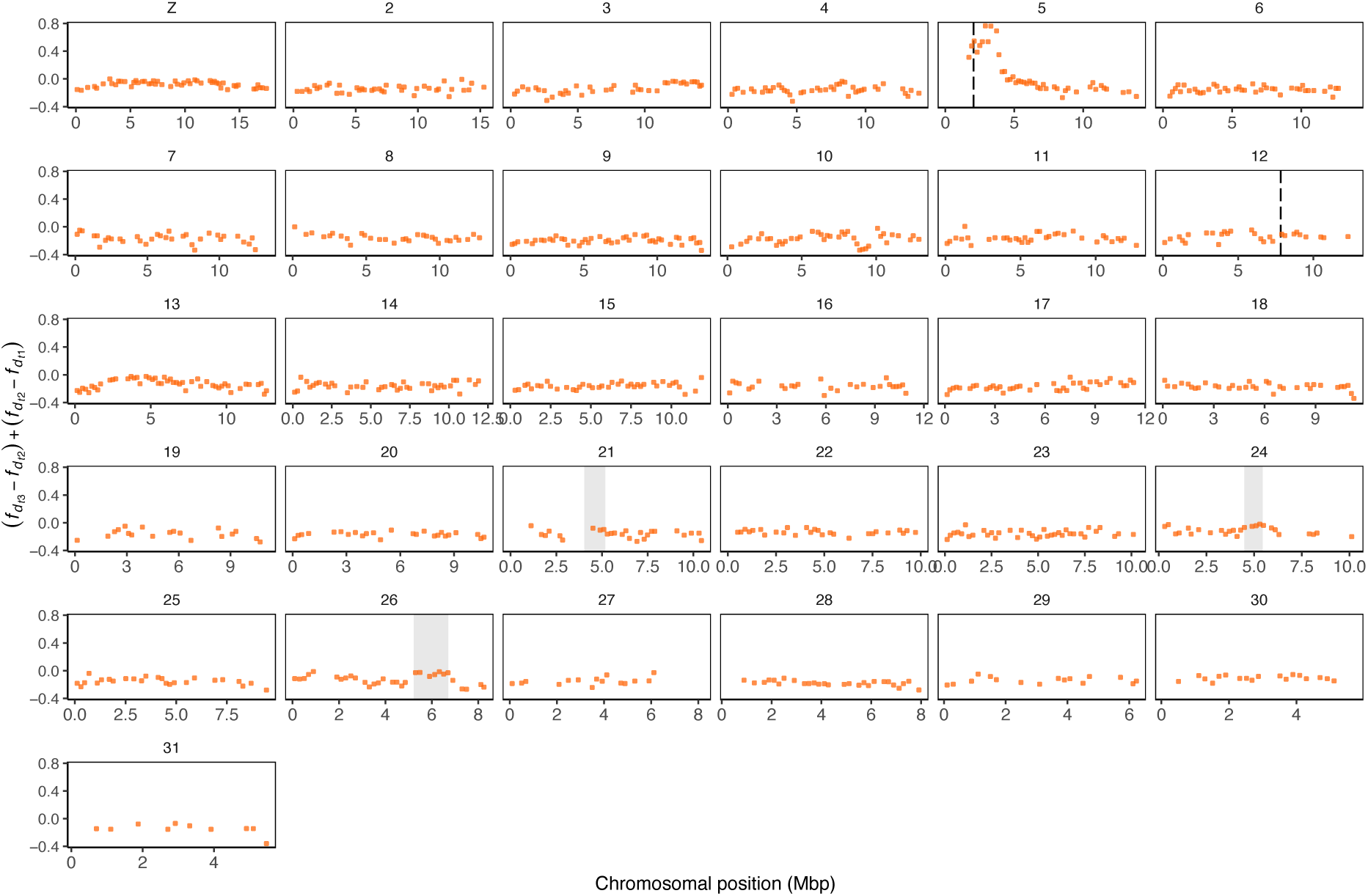
Positive change in *f_d_* over three timepoints, calculated in 200kbp windows with at least 500 usable sites per window for the topology (((ZEA, ZEA_BRAZIL), ARM)), PUN), *i.e.* gene flow from *H. armigera* to *H. zea*.

**Extended Data Figure 23:**
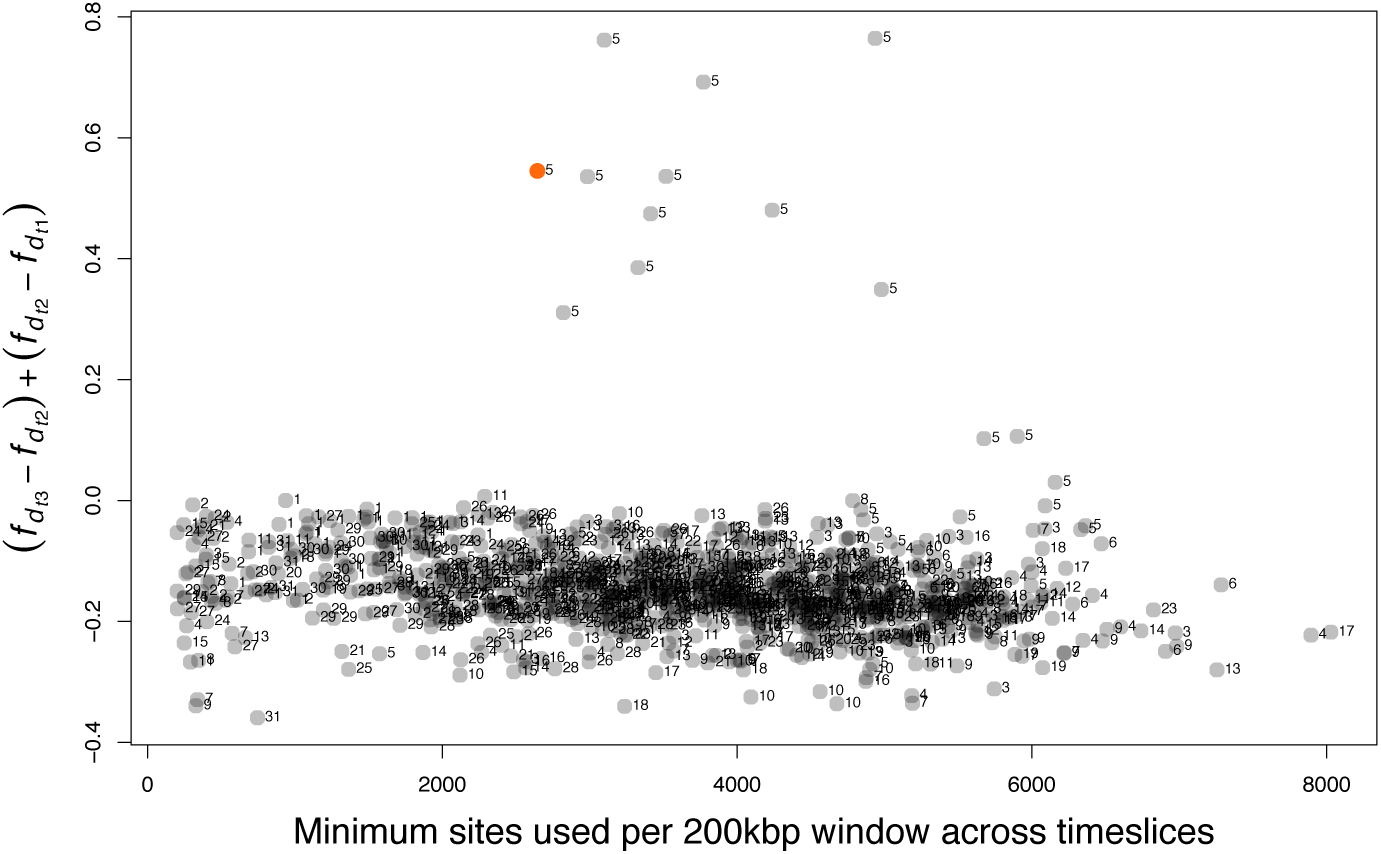
Vertical axis: Positive change in *f_d_* over three timepoints, calculated in 20kbp windows with at least 200 usable sites per window for the topology (((ZEA, ZEA_BRAZIL), ARM)), PUN), *i.e.* gene flow from *H. armigera* to *H. zea*. Horizontal axis: minimum number of sites used per window. Points are labelled with the chromosome on which the window occurs. The window on which *CYP337B3* occurs is highlighted in orange.

**Extended Data Figure 24:**
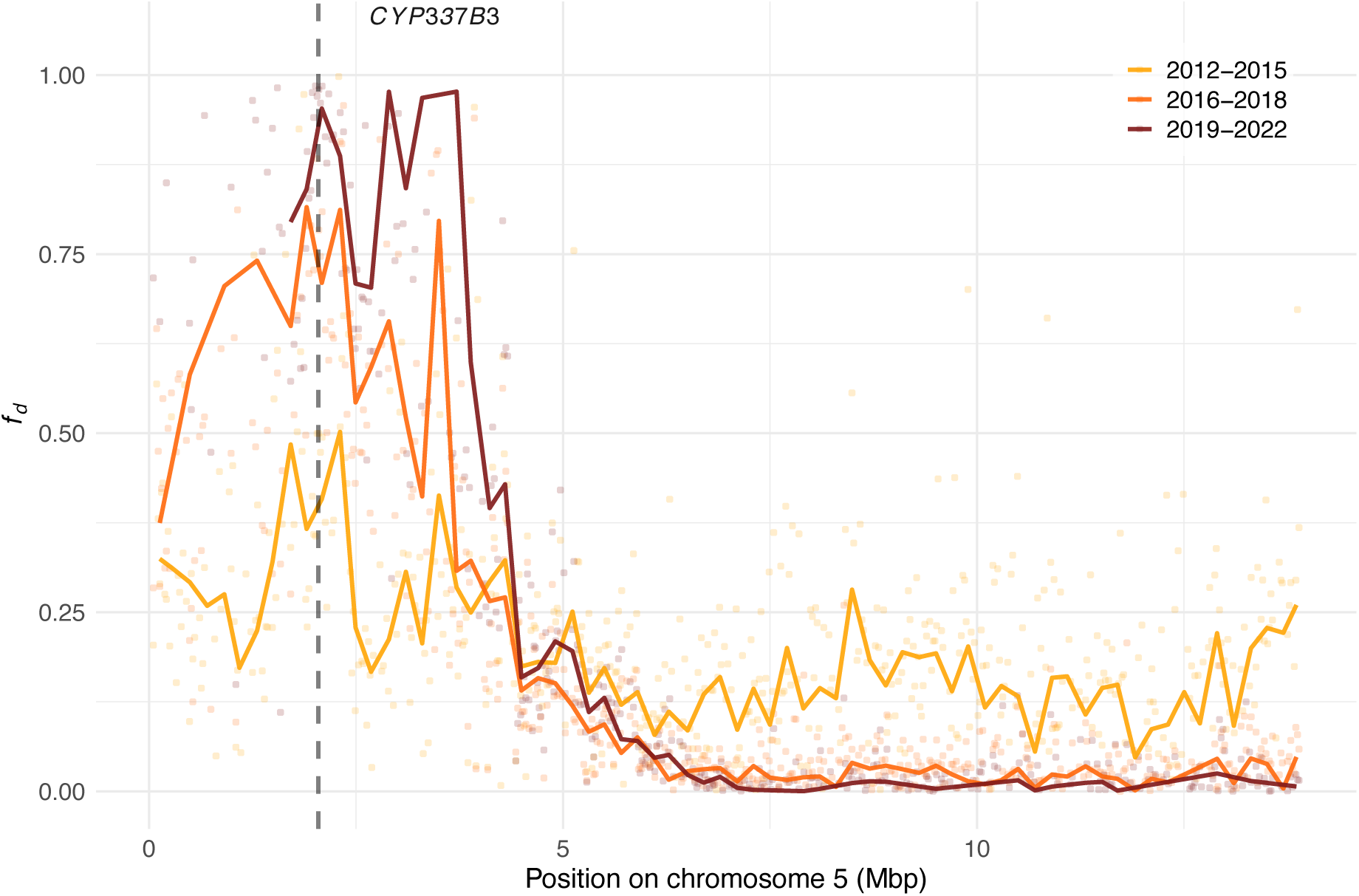
*f_d_* using the topology (((ZEA, ZEA_BRAZIL), ARM)), PUN), indicative of introgression from *H. armigera* to *H. zea*, shown for chromosome 5 for individuals split into three temporal sets. 200kbp windows with at least 500 usable sites shown as lines; 20kbp windows with at least 200 usable sites shown as points. Dasshed line indicates the position of *CYP337B3*.

**Extended Data Figure 25:**
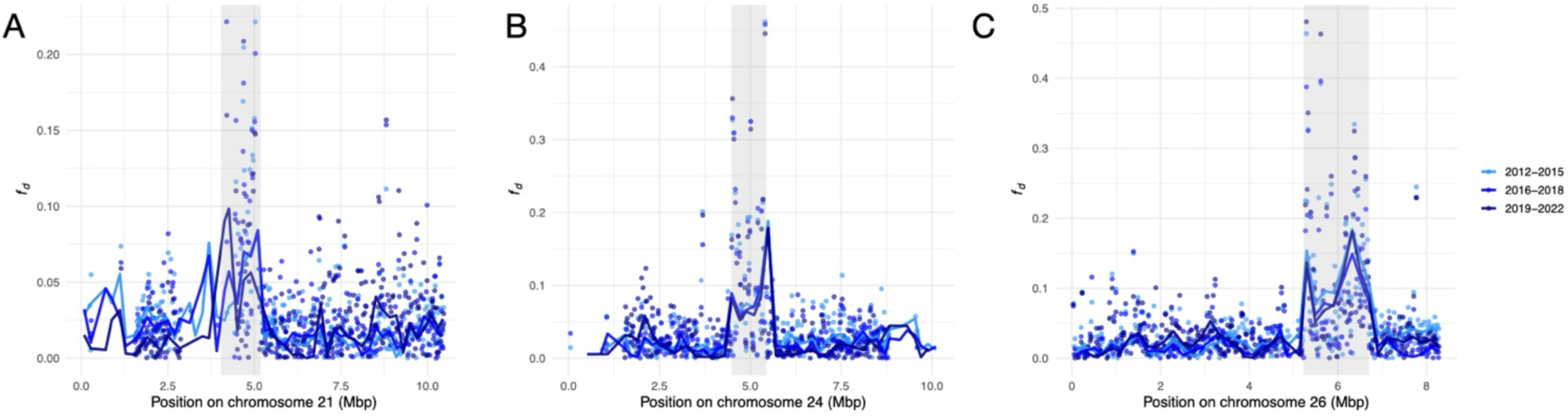
*f_d_* using the topology (((ARM, ARM_BRAZIL), ZEA)), PUN), indicative of introgression from *H. zea* to *H. armigera*, shown for chromosomes 21, 24, and 26 (A-C respectively), for individuals split into three temporal sets. *f_d_* calculated in 200kbp windows (lines) and 20kbp windows (points), with inversion breakpoints inferred based on synteny comparisons displayed in grey. There is no apparent change in the introgression signal over time in the inversion regions.

**Extended Data Figure 26:**
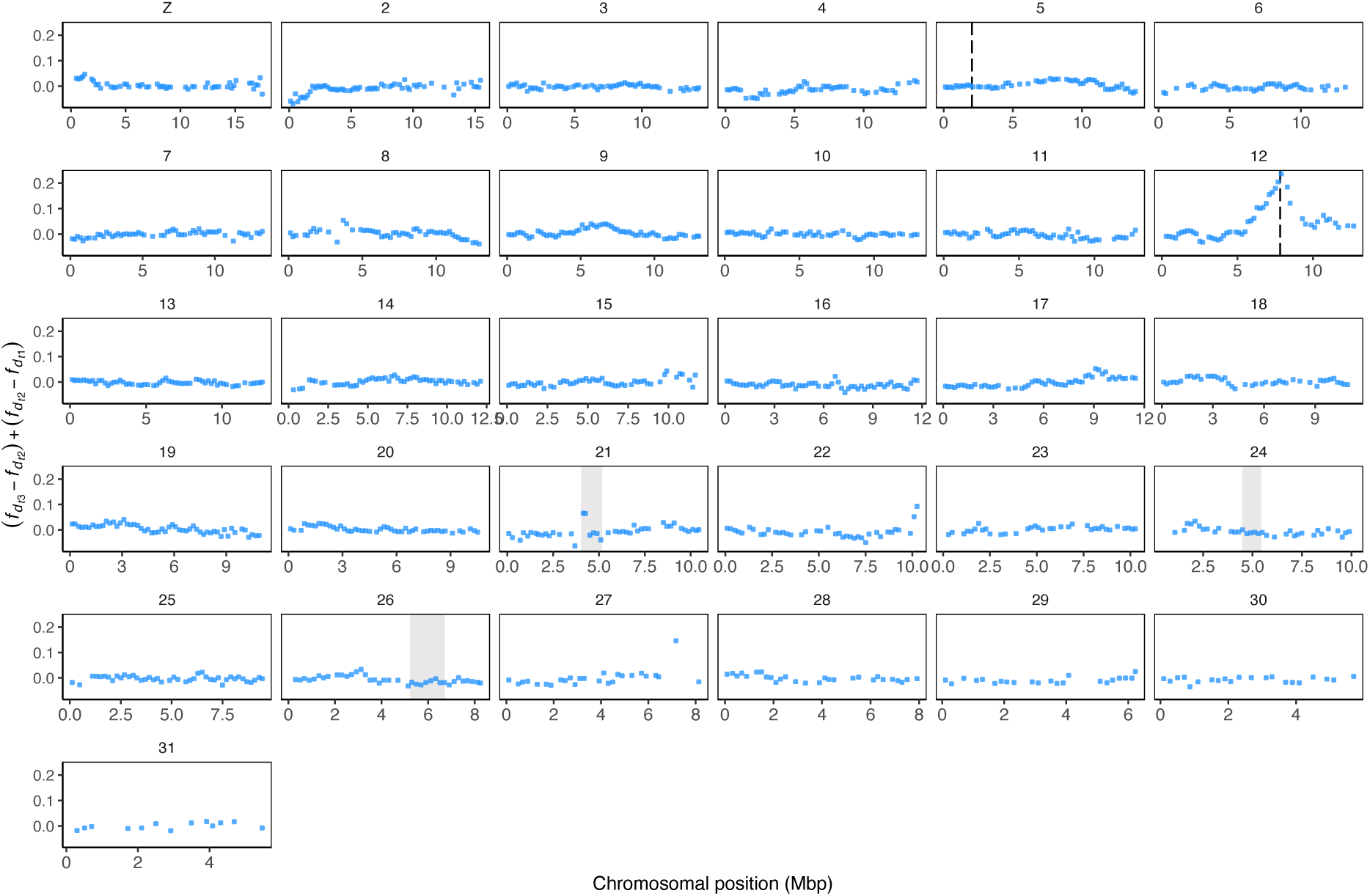
Positive change in *f_d_* over three timepoints, calculated in 200kbp windows with at least 500 usable sites per window for the topology (((ARM, ARM_BRAZIL), ZEA)), PUN), *i.e.* gene flow from *H. zea* to *H. armigera*.

**Extended Data Figure 27:**
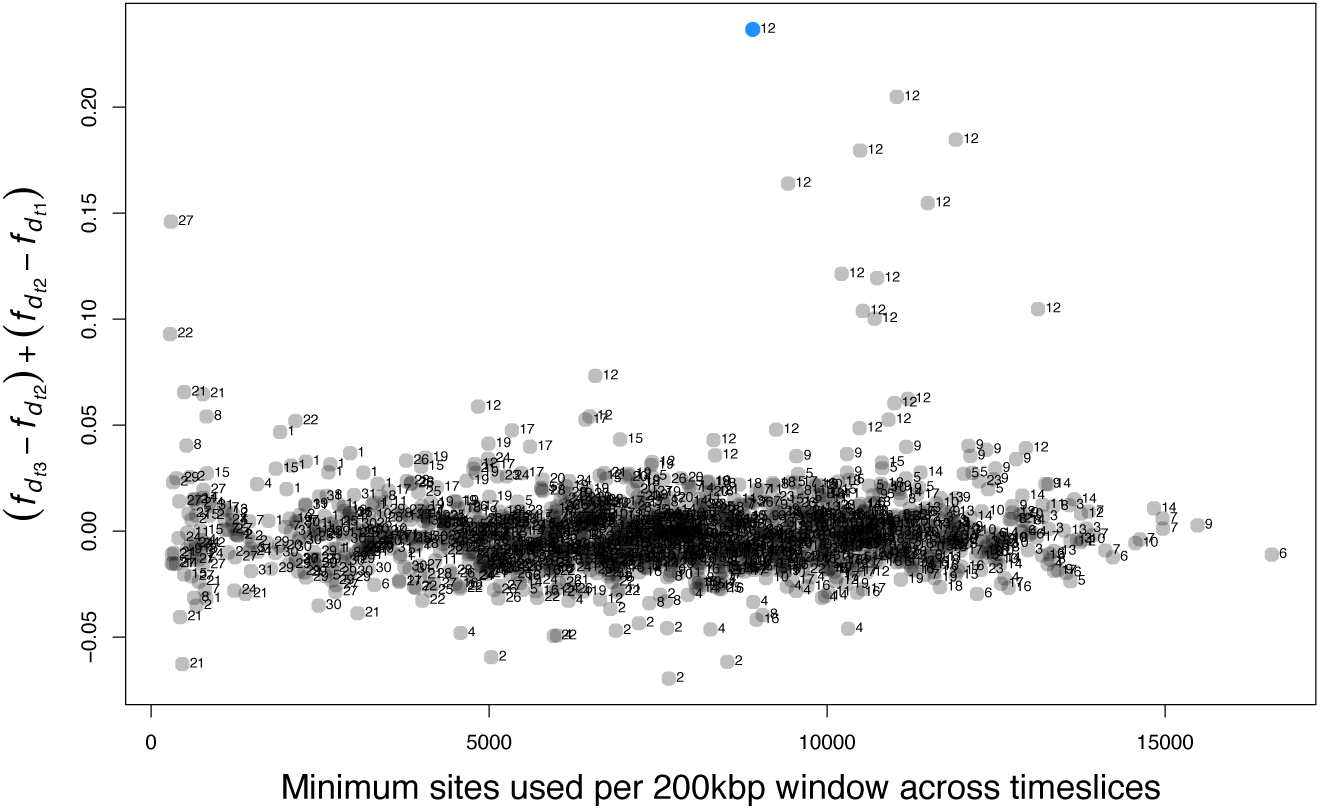
Vertical axis: Positive change in *f_d_* over three timepoints, calculated in 200kbp windows with at least 500 usable sites per window for the topology (((ARM, ARM_BRAZIL), ZEA)), PUN), *i.e.* gene flow from *H. zea* to *H. armigera*. Horizontal axis: minimum number of sites used per window. Points are labelled with the chromosome on which the window occurs. The window on which the trypsin allele cluster and the QTL for Bt resistance occurs is highlighted in blue.

**Extended Data Figure 28:**
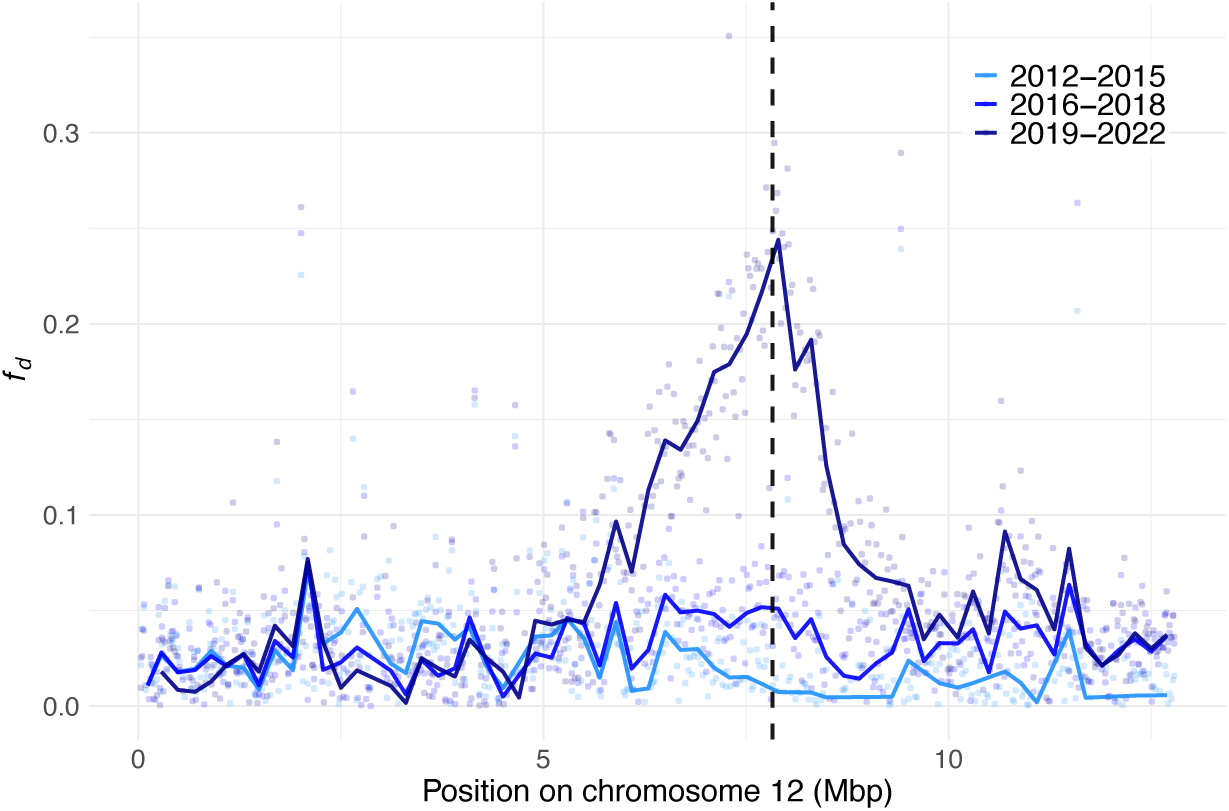
*f_d_* using the topology (((ARM, ARM_BRAZIL), ZEA)), PUN), indicative of introgression from *H. zea* to *H. armigera*, shown for chromosome 12 for individuals split into three temporal sets. 200kbp windows with at least 500 usable sites shown as lines; 20kbp windows with at least 200 usable sites shown as points. Dashed line indicates the position of the trypsin cluster.

**Extended Data Figure 29:**
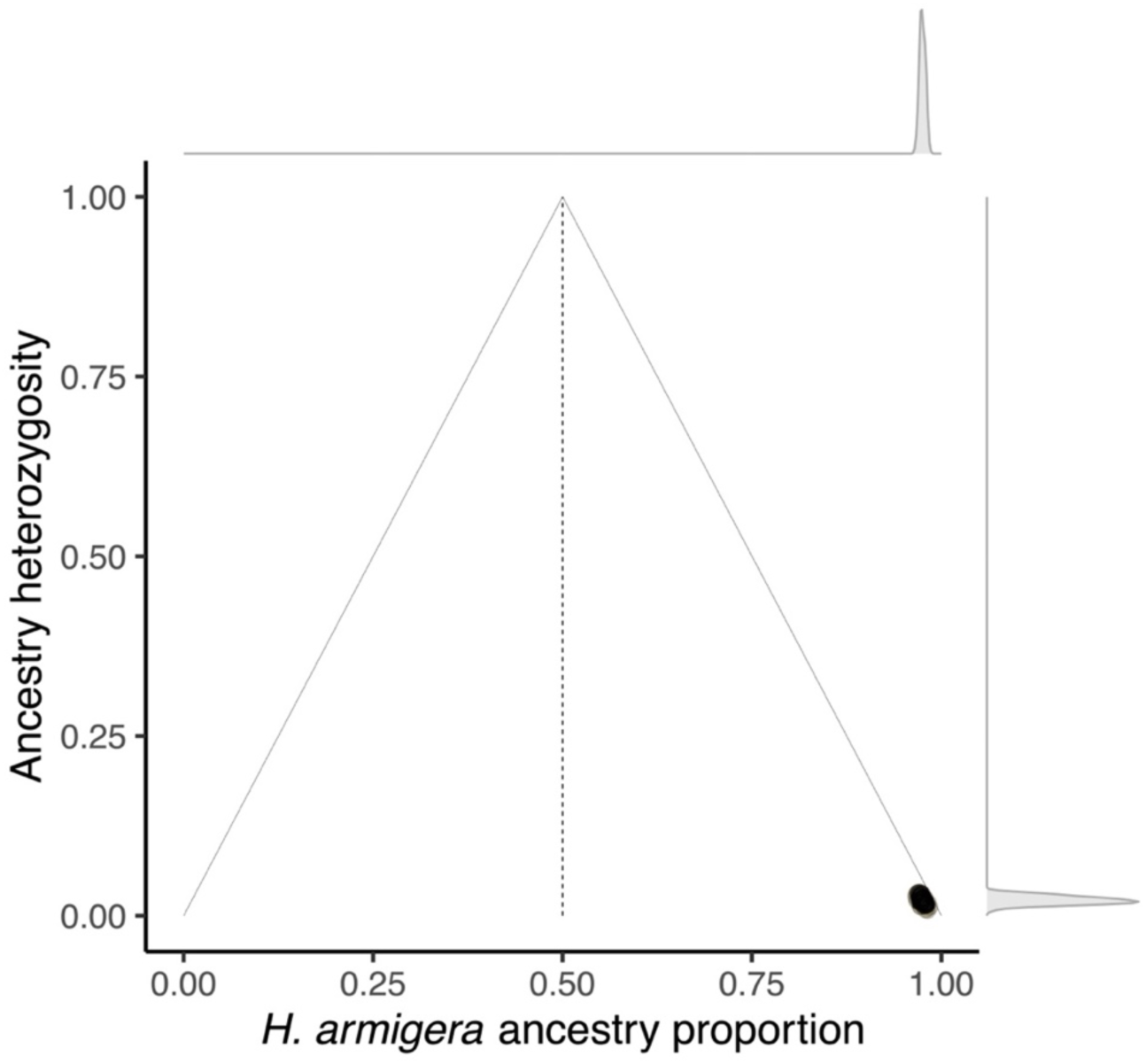
Ancestry proportion and heterozygosity (as for Figure 1C) of QTL samples using Call Set 2; all individuals are *H. armigera* (minimum *H.* armigera ancestry proportion was 0.966, mean = 0.975).

**Extended Data Figure 30:**
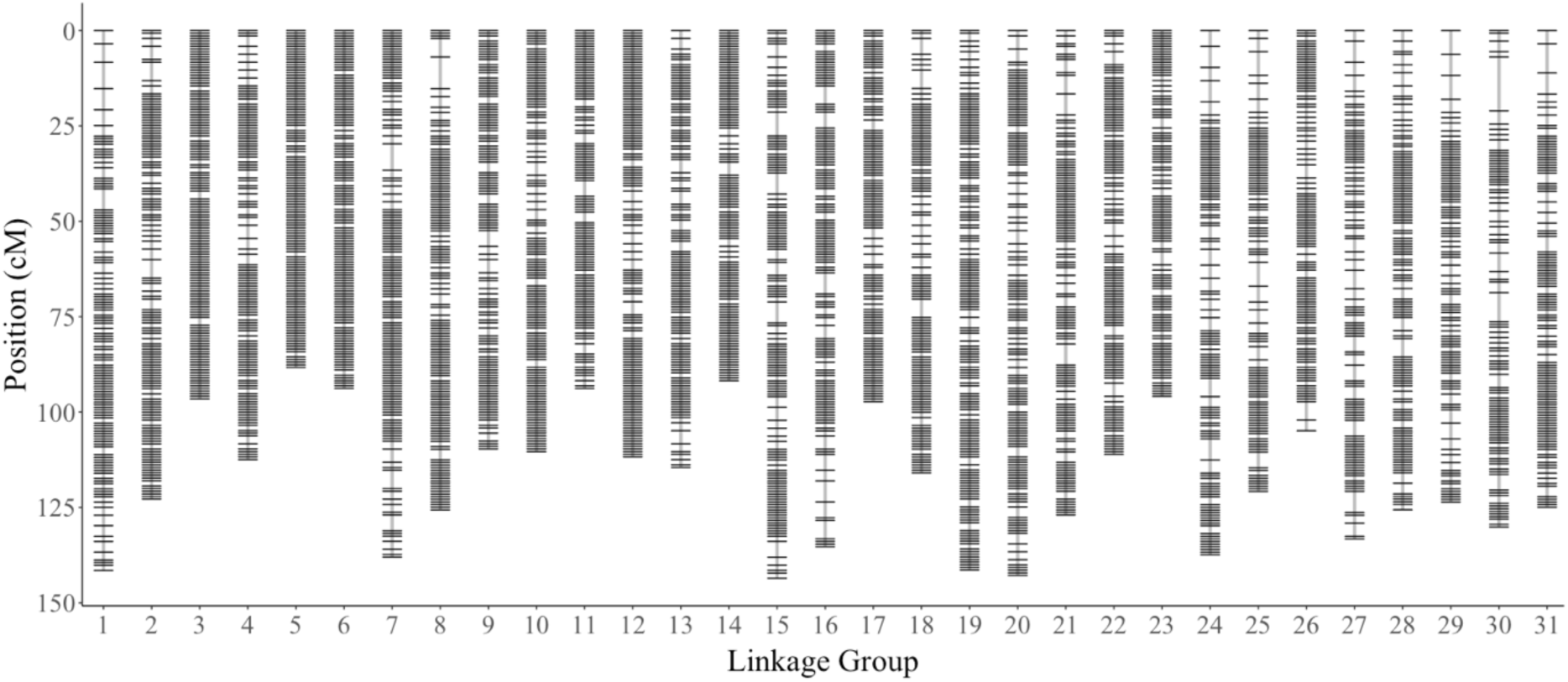
Linkage map of F2 the intercross population, derived from the cross between the *Helicoverpa armigera* susceptible and the Cry1Ac resistant strains. Linkage groups are ordered along the horizontal axis. Each line in a linkage group denotes a marker, and the vertical axis shows the genetic position of markers in cM.

**Extended Data Figure 31:**
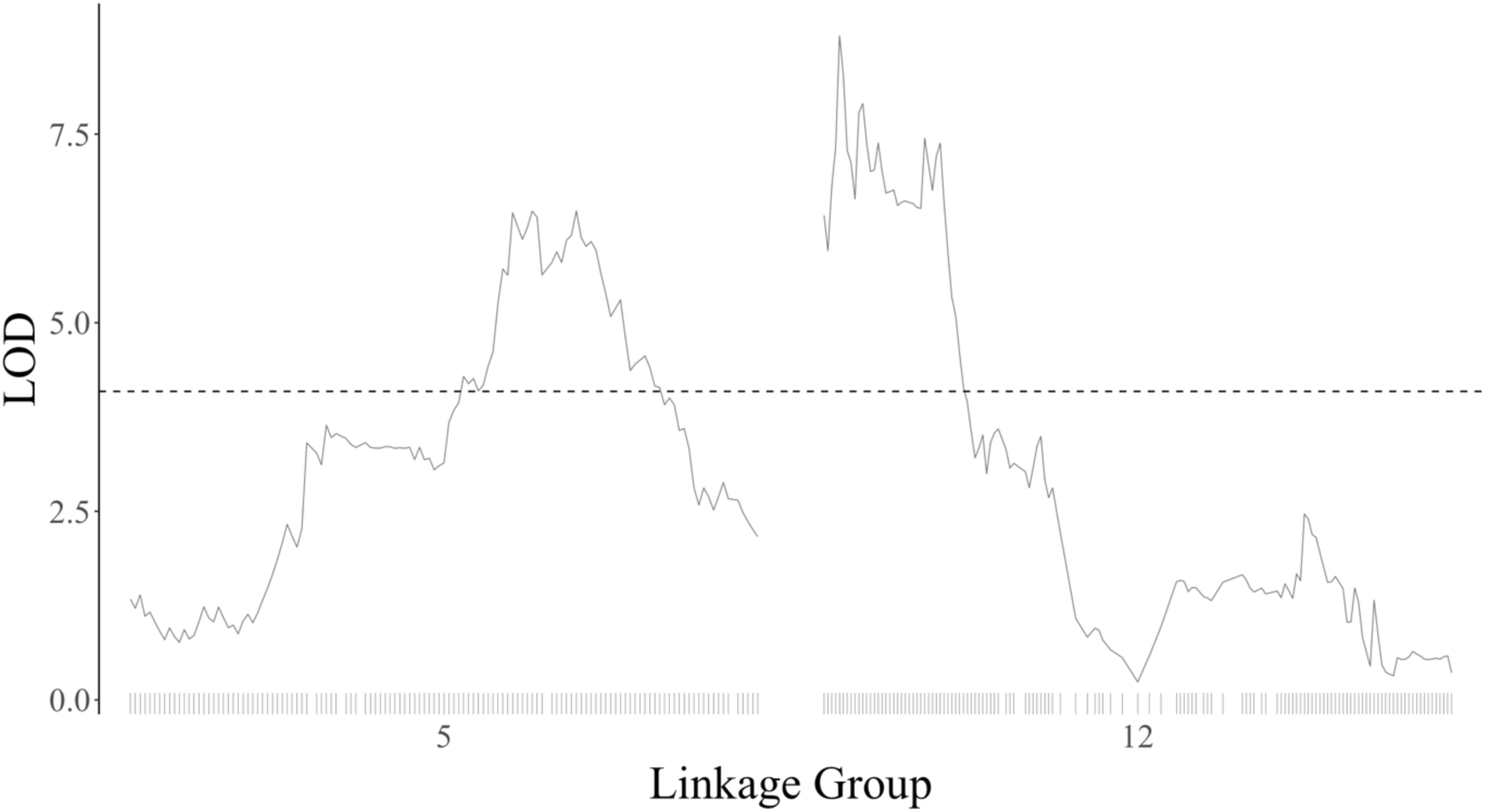
The LOD profiles of linkage groups 5 and 12, where significant QTLs for the survival trait were identified. The Y-axis shows the LOD ratio, and the X-axis shows the linkage groups 5 and 12. The lines at the bottom of the linkage group denote markers.

**Extended Data Figure 32:**
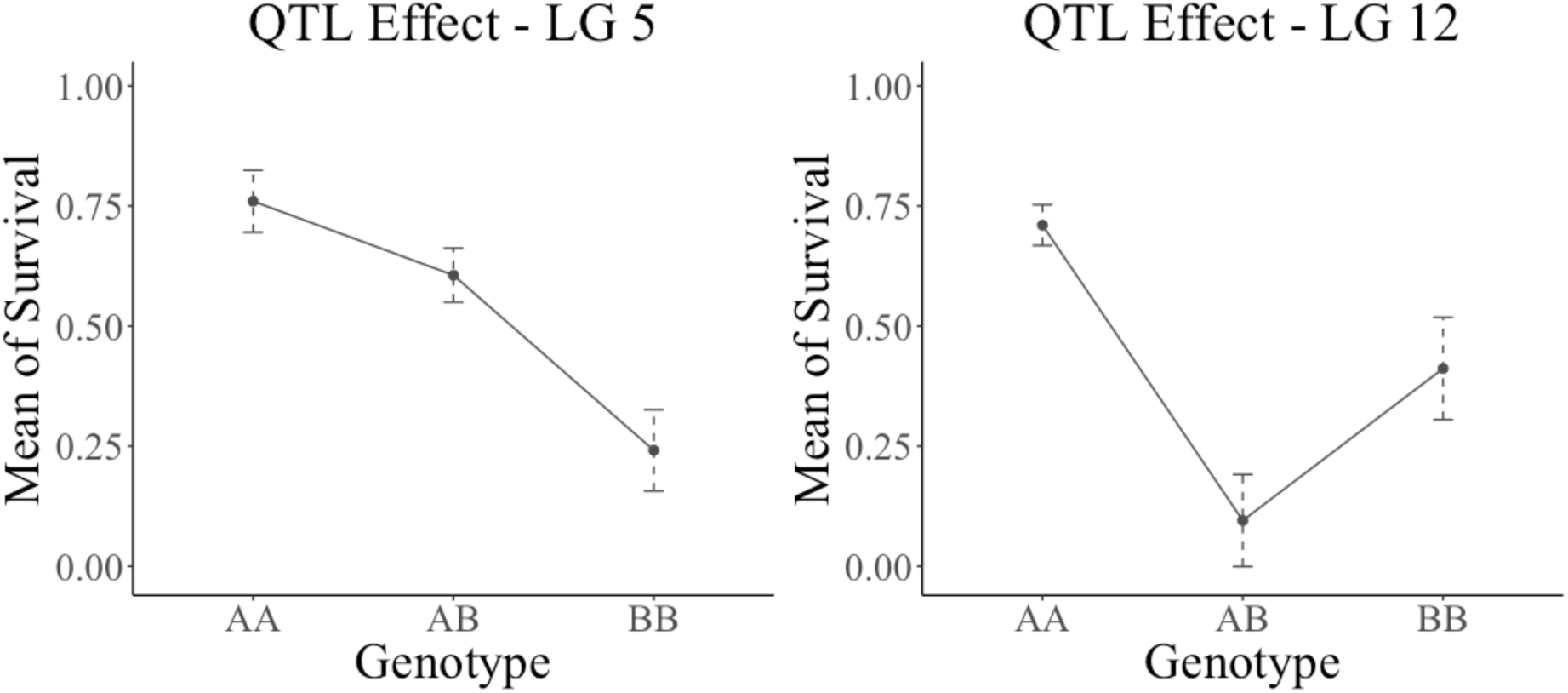
Efect plot of the main QTL on linkage groups 5 and 12. The X-axis represents the genotypes homozygote resistant (AA), heterozygote (AB) and homozygote susceptible (BB). The Y-axis represents the phenotypic value (survival ratio) after a transformation using the logit function.

**Extended Data Figure 33:**
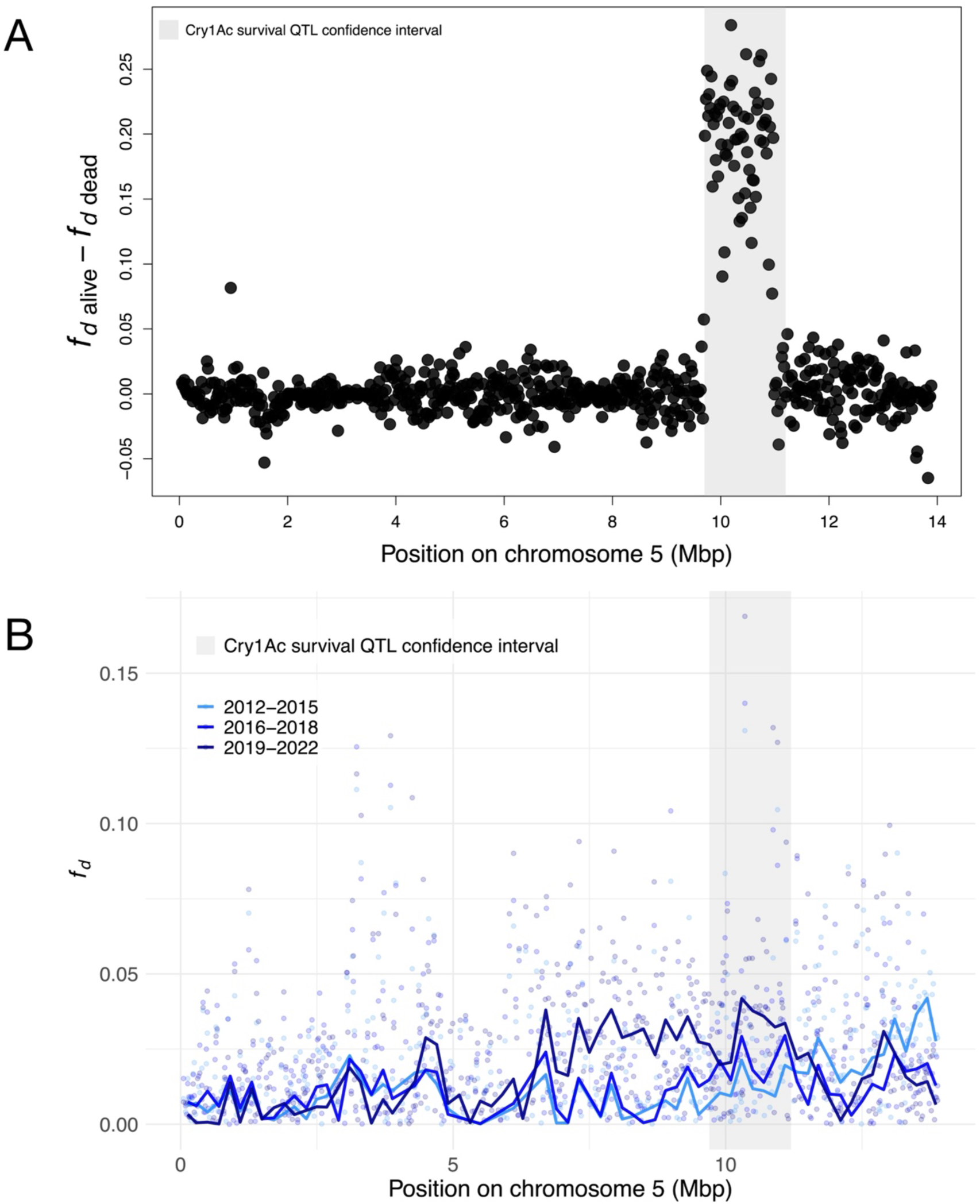
**A:** Difference in the signal of *H. zea* allele sharing between alive and dead individuals. In both cases *f_d_* was calculated using the topology (((ARM, QTL_SAMPLE), ZEA)), PUN). Grey shaded regions indicate the QTL confidence intervals. **B:** introgression from *H. zea* to *H. armigera* among wild samples divided into 3 timepoints, using the same test topology. Lines indicate 200kbp windows; points indicate 20kbp windows with at least 200 usable sites.

